# Organisational principles of long non-coding RNAs revealed by exon deletion

**DOI:** 10.64898/2026.06.17.732811

**Authors:** Sarang Bhutada, Hugo A. Guillen-Ramirez, Tina Uroda, Ines Jover Bravo, Michela Coan, Toni Hermoso Pulido, Artem Baranovskii, Annalisa Marsico, Rory Johnson

## Abstract

Long non-coding RNAs (lncRNAs) regulate cell phenotypes in health and disease, yet how function is encoded in their sequence remains poorly understood. Current models propose a modular architecture composed of discrete functional elements, but this is based on a limited set of paradigmatic examples and methods for mapping function to sequence are limited in scope and resolution. Here, we establish a high-throughput CRISPR-Cas9 strategy for dissecting lncRNA functional architecture at exon resolution. Using cell fitness as a phenotypic readout, we screened 358 exons from 107 lncRNAs across four human cell lines. We report that (1) a large proportion of exons have no detectable function, (2) a minority of exons are functional in any given cell line (19–111 exons), equivalent to one-fifth of total transcript nucleotides on average, and (3) functionality is enriched towards the 5’ end of the transcript. We developed a database of putative lncRNA functional elements, ElementaLdb, and demonstrated through statistical and experimental analyses that lncRNA function depends on transposable elements, microRNA response elements and RNA binding protein sites. These sub-genic functional maps expand the catalogue of experimentally defined lncRNA functional elements by an order of magnitude, illuminate molecular mechanisms and broadly support a modular organisation for lncRNAs.

## Introduction

Over one thousand long noncoding RNAs (lncRNAs) have been implicated in human cell phenotypes across healthy and diseased states^1^. Despite their prevalence, the mechanistic basis for this bioactivity remains poorly understood. Much of our current knowledge derives from studies of a narrow set of highly expressed lncRNAs like NEAT1^2^ and XIST^3^, which may not be representative of the broader class. Consequently, two fundamental questions remain unanswered: how is functional sequence organised within lncRNA transcripts? and how does sequence govern functionality?

A widely discussed organisational model posits that lncRNAs are modular; that functional regions are dispersed throughout the transcript and separated by flexible, non-functional spacers^4^ (Fig 1A). This ‘beads-on-a-string’ model draws support from prominent examples such as XIST, which harbours distinct domains for transcriptional silencing and chromosomal localisation that can be independently deleted without reciprocal loss of function^5^. The growing numbers of lncRNAs containing functional sub-sequences, which recapitulate partial or full activity of the parent transcript^2,6–9^ when expressed in isolation, provide further evidence for modularity and raises the possibility that the remaining sequence is dispensable. On the other hand, recent structural studies of the MEG3 lncRNA identified a long-range kissing loop interaction essential for function^10^, which is suggestive of a global folded structure akin to proteins. Whether lncRNAs are better described using the ‘beads-on-a-string’ or ‘global-fold’ model, remains an open question in RNA biology^11^.

**Figure 1:**
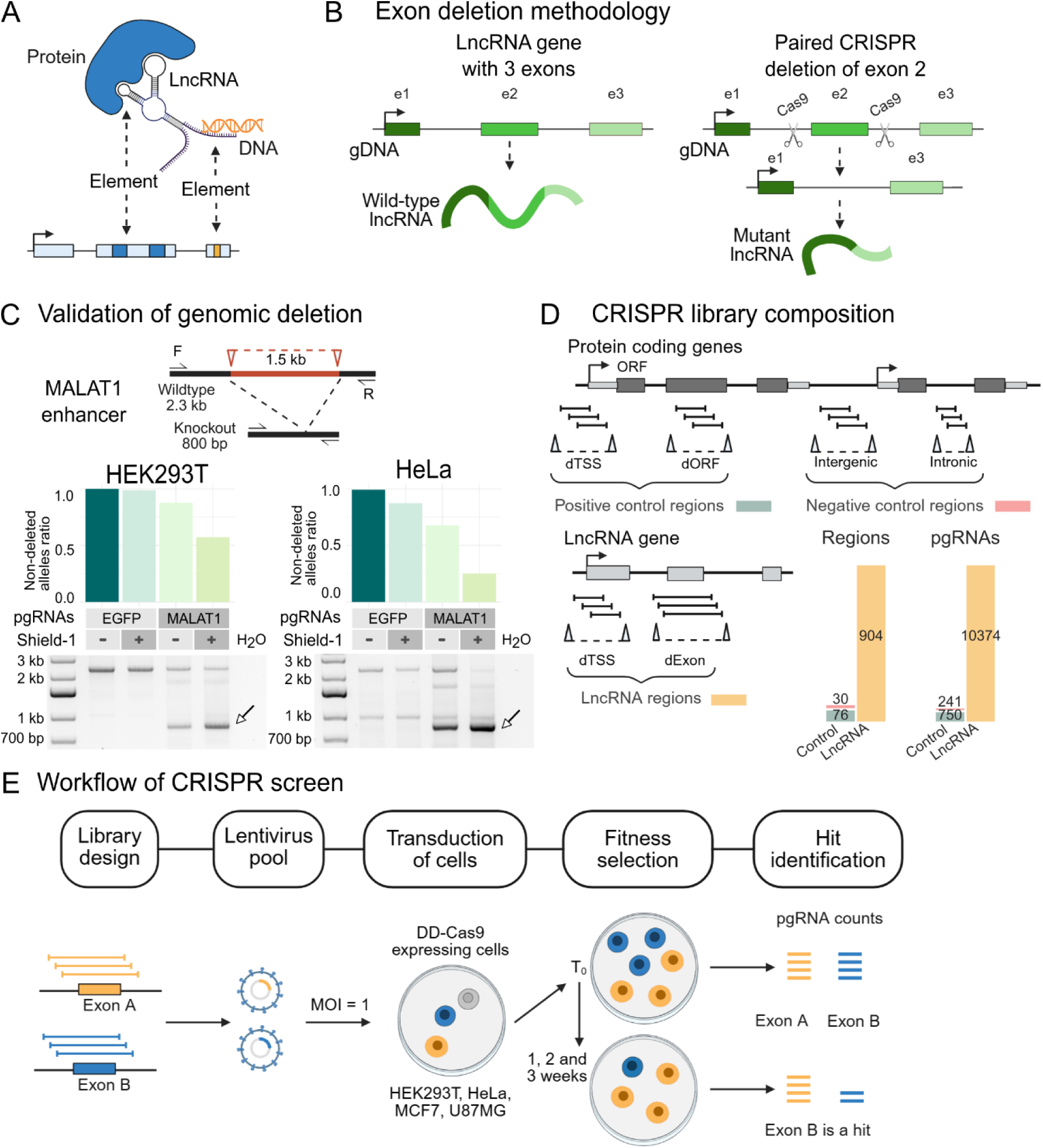
High-resolution CRISPR screen to study lncRNA exons. (A) A modular lncRNA showing protein (blue) and DNA (orange) interaction elements. (B) Schematic representation of the methodology used for exon deletion. Left: an example lncRNA with 3 exons (wild-type lncRNA). Right: paired CRISPR deletion targeting the second exon (e2). pgRNAs flanking e2 are used to excise the target DNA resulting in a mutant lncRNA. (C) Validation of genomic deletion using inducible Cas9. Non-deleted alleles ratio was quantified by PCR using pgRNAs targeting the MALAT1 enhancer, with and without Shield-1 treatment in HEK293T and HeLa cell lines. EGFP pgRNAs were used as non-targeting control. The bar plot shows mean of two technical replicates. The gel electrophoresis of the wild-type and genomic deletion amplicon of MALAT1 enhancer is shown below. F: forward primer, R: reverse primer. The arrows indicate the DNA bands with the expected deletion. (D) Design of the CRISPR library. Top: pgRNAs targeting TSS (dTSS) and ORF (dORF) of protein coding genes (PCG) were used as positive controls, while introns of PCG and intergenic regions were negative controls. Bottom: pgRNAs were designed to target TSS (dTSS) and exons (dExon) of lncRNAs. The bar plot shows the size of the library in terms of total number of regions targeted in each category and the corresponding number of pgRNAs. (E) Workflow of the CRISPR screen. The pgRNA library was transduced (MOI=1) in four DD-Cas9 expressing cell lines. Cells were selected with puromycin and cultured for 1, 2 and 3 weeks. At all the timepoints, including T0, DNA was isolated from harvested samples to sequence the pgRNAs, followed by statistical analysis to identify the hits.

The mechanistic basis for lncRNA activity can be attributed to interactions with other biomolecules, namely protein, RNA and DNA (Figure 1A). RNA binding proteins (RBPs) recognise RNAs through short sequence motifs, as mapped experimentally by methods such as eCLIP^12^, enabling spatial targeting of proteins or assembly of multi-protein complexes^13,14^. LncRNAs can form Watson-Crick hybrids with other RNA transcripts including microRNAs^15^. They can also target specific genomic loci through formation of RNA-DNA hybrids such as R-loops^16^. Transposable and repetitive elements have also been shown to serve as functional domains within lncRNAs, potentially through pre-formed protein-binding motifs^17,18^. Evolutionary conservation offers an orthogonal lens for identifying functional sequence^19,20^, though it does not, *a priori*, reveal the underlying mechanism.

Nonetheless, these insights rest on a handful of case studies, and the extent to which they generalise remains unknown. The methods used to interrogate individual lncRNAs, including overexpression of truncated constructs^21^, deletion by CRISPR^2^, are laborious and require bespoke experimental design for each transcript. A systematic, high-throughput assessment of lncRNA sequence function at sub-genic resolution is lacking.

To address this gap, we developed a high-throughput, dual-guide CRISPR deletion strategy to reveal the landscape of functional exons in a large panel of lncRNAs across four human cell types, enabling us to shed light on the features and functional elements that underwrite lncRNA bioactivity.

## Results

### A high-throughput dual CRISPR deletion method to identify functional lncRNA exons

To map functional nucleotides within lncRNAs at sub-genic resolution, we designed a high-throughput, CRISPR-Cas9 approach that quantifies the function of individual exons. This methodology uses paired guide RNAs (pgRNAs) to delete exons together with their splice signals, resulting in a mutant transcript that lacks the targeted exon (Fig 1B). As a functional readout, we chose cell fitness, as measured after serial passaging, similar to previous gene-level screens^22,23^.

To achieve precise temporal control over DNA deletion and avoid deleterious effects of constitutive Cas9 expression^24^, we fused it to a destabilizing domain (DD)^25^ to engineer an inducible Cas9, enabling activation of CRISPR deletion upon addition of Shield-1 reagent to the culture medium. This inducible system was validated by targeting an enhancer within the MALAT1 lncRNA locus, which effectively deleted the intervening DNA, whereas control pgRNAs had no effect (Fig 1C and S1A). Shield-1 treatment enhanced deletion efficiency as expected, although some background deletion was observed in its absence. Overall, the inducible approach yielded specific and efficient removal of targeted genomic DNA.

We designed a pgRNA library targeting exons of lncRNAs with previously reported roles in cell fitness (Fig 1D). High-confidence fitness-promoting lncRNAs were curated from the Cancer LncRNA Census 2^26^ and a published CRISPRi screen^22^. To achieve sensitivity and overcome inefficient editing, each exon was targeted by 10 distinct pgRNAs (Fig S1B) with a median deletion length of ∼1 kb (Fig S1C). The library also included extensive controls (Fig. S1D): (a) 20 regions targeting open reading frames (ORFs) of essential protein-coding genes (‘dORF’, positive control for screen performance); (b) 10 regions targeting transcription start sites (TSSs) of essential genes, (‘dTSS’, positive control for deletion efficiency, since both the double-stranded breaks are required for TSS deletion resulting in gene silencing), (c) 76 non-exonic negative control regions with no expected phenotypic impact.

With this library, we screened four well-studied human cell lines: HEK293T (non-cancerous), HeLa (cervical cancer), MCF7 (breast cancer) and U87MG (glioblastoma). The screen followed a dropout format (Fig 1E), in which ‘hits’ (also referred to as ‘fitness-promoting’ exons) are defined by depletion of their pgRNAs over time (Exon B), while ‘non-hits’ (also referred to as ‘fitness-neutral’ exons, i.e. not required for cell fitness under these conditions) maintain stable pgRNA representation (Exon A). The quality of screen results was supported by favourable pgRNA uniformity before (Fig S1E, F) and after transduction (Fig S1G) as well as correlation between biological replicates (Fig S2A). The area under the ROC curve (AUROC) based on the controls was >0.8 for most cells and timepoints (Fig S2B), underlining the robustness of the screen.

Positive controls further supported screen sensitivity: 66% of positive control regions were correctly identified in the majority of cell lines (Fig 2A). dTSS, the most relevant benchmark, showed a sensitivity of 90% (Fig S2C). Maximum sensitivity was achieved at the 3-week timepoint (Fig S2D). We defined exon hits from two timepoints as ‘high-confidence’ and focused on these for all downstream analyses. Among 38 negative controls, only one was called as a hit (in HeLa), confirming high specificity of the screen (Fig S2E).

**Figure 2:**
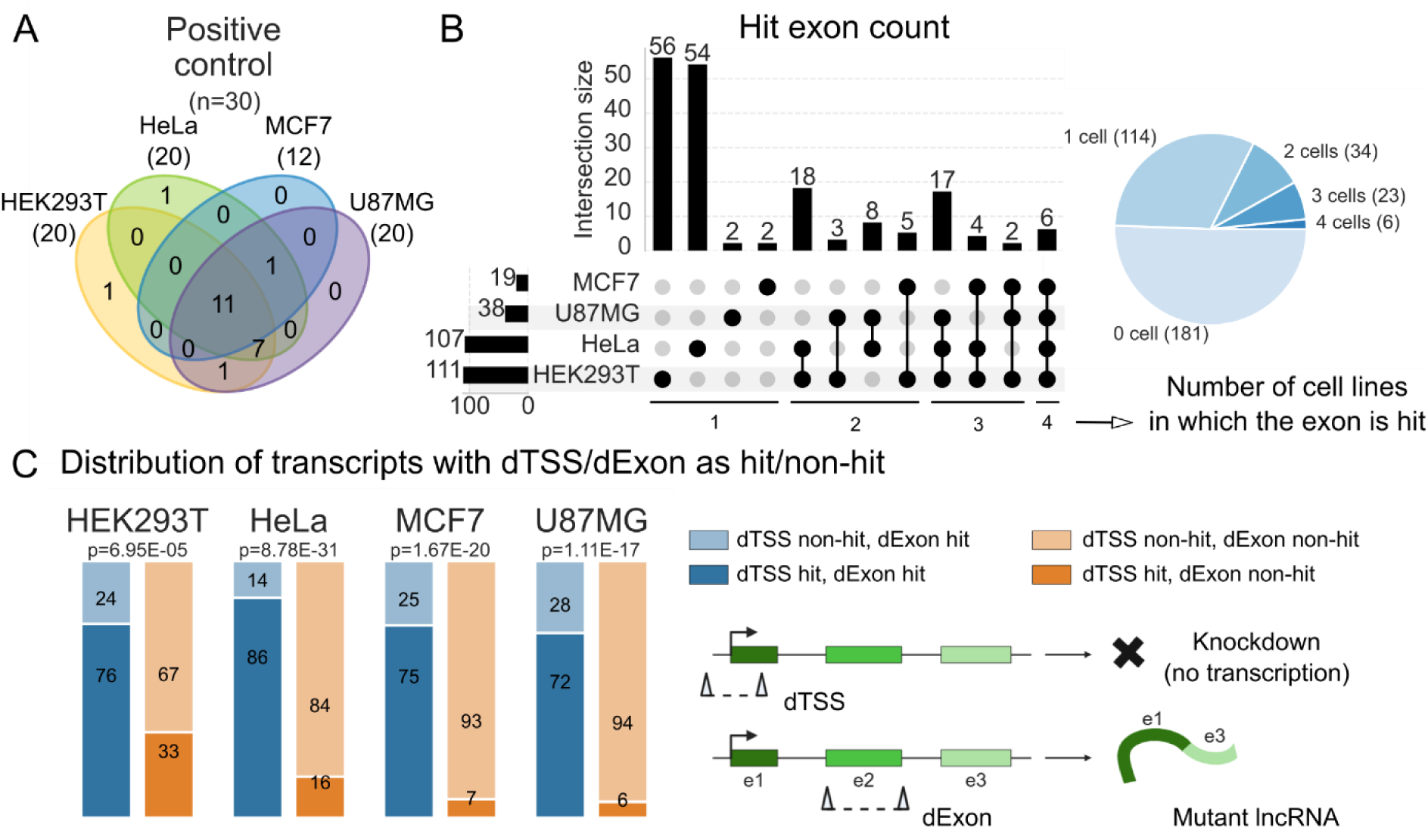
Results of the paired CRISPR screen. (A) Number of positive controls identified as high-confidence hits in each cell line. The numbers in the brackets are the total number of positive controls identified in each cell line out of total 30 controls. (B) Number of exons identified as hits from the CRISPR screen. The vertical bars show the number of hits that are cell line-specific and common among different cell line combinations as indicated by the dot matrix below. The horizontal bars show the total number of hits in each cell line. The pie chart shows the total number of hits common among 0, 1, 2, 3 or 4 cell lines. (C) Proportion of hit transcripts in two populations of transcripts: transcripts which have at least one internal hit exon (blue) and transcripts which do not have a hit exon (orange). The p values were calculated using the Chi-Square test.

Out of the 358 targeted lncRNA exons, 111, 107, 19 and 38 were identified as fitness-promoting in HEK293T, HeLa, MCF7 and U87MG respectively (Fig 2B). HEK293T and HeLa had a higher number of both total and cell line specific hits compared to MCF7 and U87MG. In contrast to positive controls, the majority of lncRNA exons displayed cell line-specific activity (i.e. hit only in a subset of cell lines) (Fig 2B). Among all the lncRNAs having at least one hit exon, 86% contained at least one exon with cell line-specific activity.

In addition to internal exon deletions, the library also targeted the TSS and first exon of the lncRNAs. These deletions are expected to silence the entire transcript and serve as an important readout of whole gene function. If a lncRNA contains at least one downstream hit exon, then its TSS deletion is expected to yield the same phenotype. Indeed, we observe this for every cell line (Fig 2C). The minority of lncRNAs where this is not true yield an estimated false-negative rate of 14–28%, comparable to the 10–40% rate estimated by the positive controls (dTSS of protein coding genes, Fig S2C). These false negatives likely reflect inefficient pgRNA-mediated deletion or alternative TSSs that were not targeted by the library. Together, these results establish the sensitivity and precision of the functional exon maps.

### Functional exon maps reveal sparse, context-dependent modularity

We next examined the functional exon maps in greater detail (Fig 3A). As expected, some lncRNAs displayed cell line-specific activity: for example, MIR31HG was a hit only in HeLa, with Exon 4 underwriting this effect. Strikingly, even among lncRNAs that promoted fitness in multiple cell types, the specific exons responsible differed between cell types. PVT1, for instance, promoted fitness in all four cell lines, yet via dramatically different exon combinations in each. Similarly, the fitness effect of PAX8-AS1 was mediated by Exon 3 in HEK293T but Exon 4 in HeLa.

**Figure 3:**
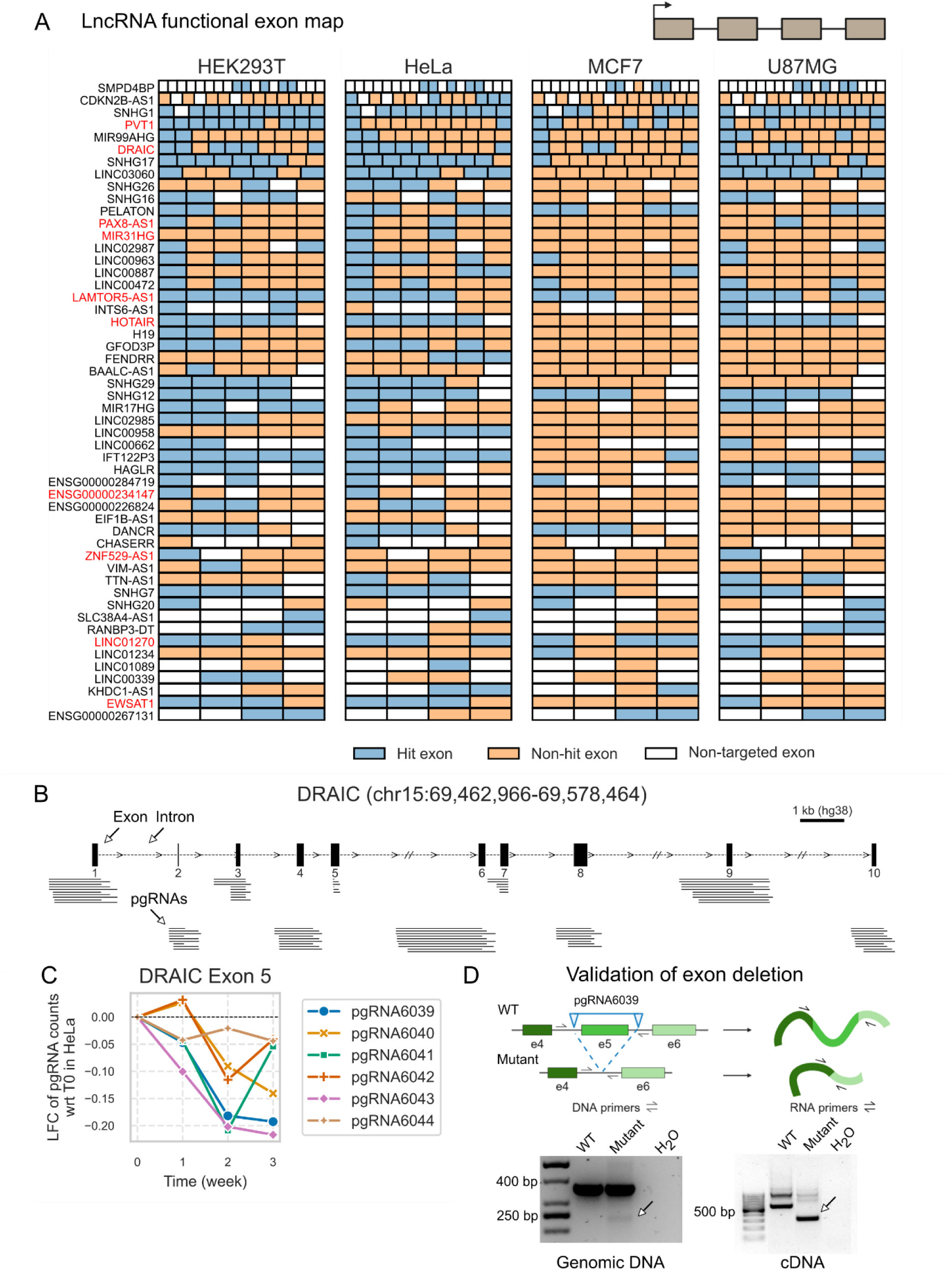
Functional landscape of lncRNA exons. (A) Distribution of exon functionality in the screened lncRNAs across the four cell lines. The transcripts are oriented left to right. Hit exons are coloured in blue, non-hit exons are coloured in orange, while non-targeted exons, which are not screened, are shown in white. Example lncRNAs, discussed in the text, are highlighted in red. (B) DRAIC transcript model showing the exons (black rectangles) and introns (dashed lines). The pgRNAs targeting the exons are shown below each of the 10 exons as lines. (C) Log2 fold change (LFC) of the pgRNA counts targeting Exon 5 of DRAIC in HeLa at three timepoints, relative to the initial timepoint T0. (D) Validation of DRAIC exon 5 deletion targeted by pgRNA6039. Top: As indicated by the semi-arrows, primers used for amplifying the genomic DNA (gDNA) were designed in the flanking introns, while primers for reverse transcription of RNA were in the neighbouring exons. Bottom: Agarose gel showing PCR products with template gDNA and cDNA from RNA template from HEK293T cells transfected with non-targeting pgRNA (WT) or pgRNA6039 (Mutant).

Another interesting feature of these maps is the sparsity of functional sequence: the majority of lncRNA exons had no detectable contribution to fitness, even when dTSS indicate that their host lncRNA is active. For instance, ENSG00000234147 and ZNF529-AS1 were hits in two of four cell lines, yet their screened internal exons appeared entirely dispensable, suggesting that all functionality is encoded within the first exon. Even after correcting for the estimated false negative rate of the screen (10–40%, based on positive controls), 65–90% of exons are fitness-neutral in any given cell line, and 37–50% showed no measurable activity across all four cell lines. Thus, lncRNA function is encoded by a minority of exonic sequence, with the bulk of the transcript tolerating deletion without phenotypic consequence, at least for cell fitness.

Among the top hits was the lncRNA ‘Downregulated RNA In Cancer’ (DRAIC), previously reported as either a tumour suppressor or oncogene depending on context^27^. The pgRNAs targeting DRAIC are shown in Fig 3B. The TSS and Exon 5 of DRAIC emerged as hits in all four cell lines (Fig 3A). pgRNAs targeting this exon showed progressive depletion across timepoints, supporting its fitness-promoting role (Fig 3C). Genomic PCR confirmed effective deletion of the target region in HeLa cells, and RT-PCR together with Sanger sequencing verified production of a truncated DRAIC transcript lacking Exon 5 (Fig 3D), supporting the premise that exon deletion leads to the expected mutant transcript.

The essential exons identified in PVT1 and HOTAIR are consistent with prior mechanistic studies: Exons 2 and 9 of PVT1 have been shown to be required for its oncogenic activity across multiple cancer types^28,29^, and an 89-nucleotide domain of HOTAIR spanning Exons 4 and 5 constitutes the minimal binding motif for the PRC2 complex^30^; all of which correctly reported by our screen (Fig 3A). Overall, these data represent the most comprehensive map of lncRNA functionality at sub-genic resolution.

### Experimental validation of exon functionality

To further validate the screen results, we tested selected exons individually using a fitness competition assay. Mutant cells lacking exons were co-cultured with wild-type (WT) cells, and their relative abundance was tracked over time by fluorescence (Fig 4A). As expected, deletion of the ORF of the essential gene RPS5 caused a dramatic loss of fitness, while targeting the negative control AAVS1 locus did not (Fig 4B). Consistent with the screen, cells lacking DRAIC Exon 5 were significantly outcompeted, whereas those lacking adjacent non-hit Exon 6 were not (Fig 4C). We could similarly validate hit and non-hit exons of LINC01806 (Fig 4D). Four additional non-hit exons from different lncRNAs were tested and, in every case, their deletion resulted in neutral or increased fitness (Fig S3A), corroborating the screen results.

**Figure 4:**
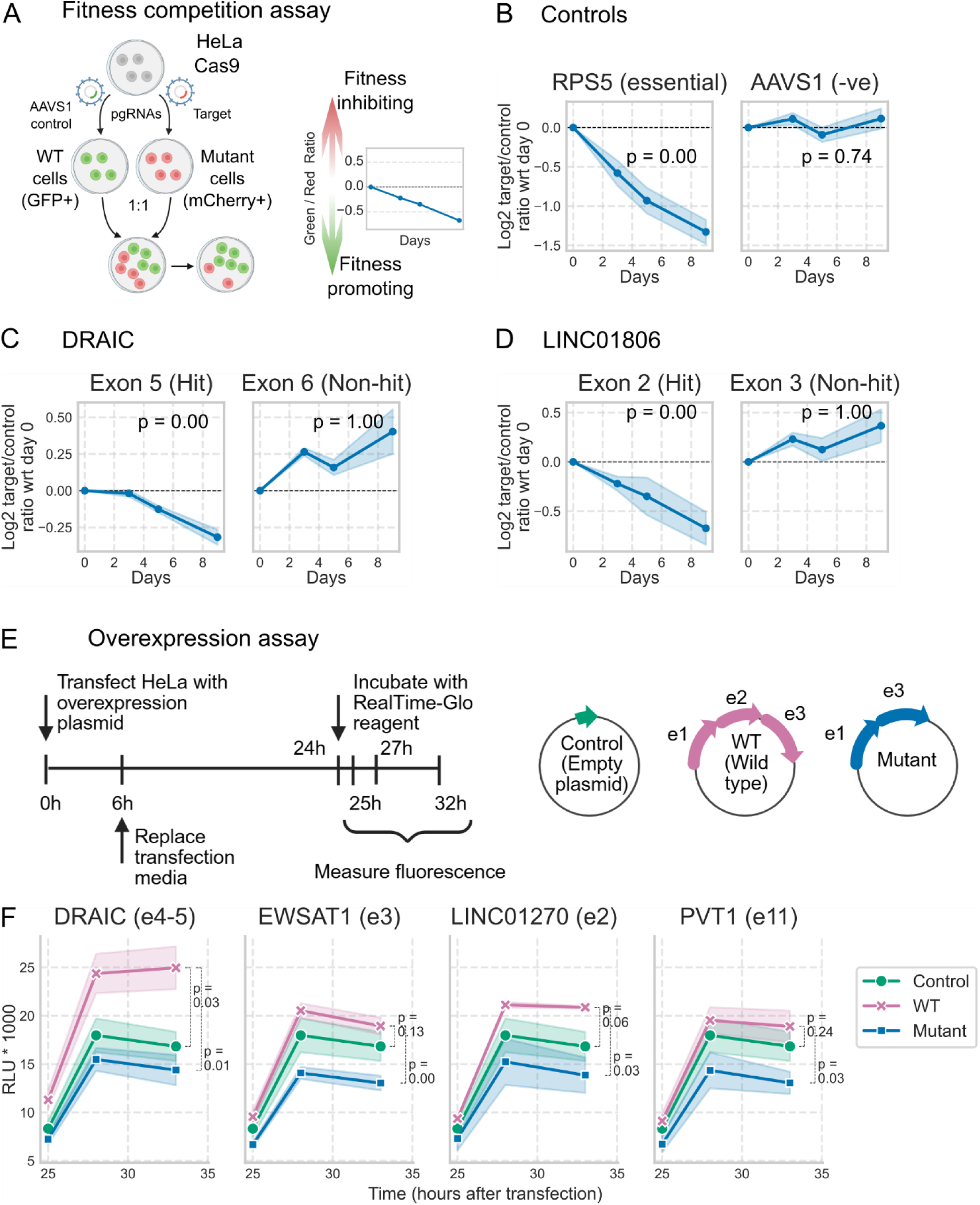
Validation of the screened exons. (A) Workflow of the fitness competition assay. HeLa cells expressing different fluorescent markers corresponding to pgRNAs for control AAVS1 locus (GFP, green) or indicated targets (mCherry, red) were measured by flow cytometry over time: 0, 3, 5, 9 days after 1:1 co-culture. The log2 transformed ratio of green and red cells normalized to T0 was plotted. (B) pgRNAs targeting the ORF of essential ribosomal protein RPS5 were used as a positive control, while another set of pgRNAs targeting AAVS1 locus were used as a negative control. pgRNAs for hit and non-hit exons in two lncRNA transcripts were tested: DRAIC (C) and LINC01806 (D). Error bars indicate standard error with at least 3 biological replicates. Statistical significance was estimated using mixed linear model. (E) Design of the overexpression assay. Cells were transfected with control, WT or mutant constructs and luminescence values were measured at 25h, 27h and 32h post transfection. (F) Growth curves of HeLa cells transfected with different plasmid constructs to overexpress the WT or mutant form for a target lncRNA. The target lncRNA gene and exon is mentioned in the title of each line plot. Error bars indicate standard error with 3 biological replicates. Statistical significance was calculated using one-sided Student’s t-test by comparing area under the growth curve.

Genomic deletion can have unintended consequences such as disruption of overlapping regulatory DNA elements, altered RNA processing, or indirect effects on neighbouring genes. Therefore, we sought to confirm exon bioactivity directly at the RNA level, providing a realistic context of the full transcript. We designed a series of expression constructs encoding either full-length wild-type (WT) or exon-deleted (Mutant) lncRNA sequences and compared their effects on cell proliferation (Fig 4E). We prioritised four lncRNAs based on activity in multiple cell lines and presence of clearly defined fitness-promoting exons: DRAIC, EWSAT1, LINC01270 and PVT1. In all cases, the results were consistent with expectation: WT lncRNA expression promoted fitness above the control plasmid, while mutant lncRNA, lacking the hit exon, abrogated this effect (Fig. 4F).

Previous studies have demonstrated that expression of individual exons can result in phenotypic effects mimicking the full-length lncRNA^7,8^. We tested this for the DRAIC exon 5 (Fig S3C) but observed no fitness effect from single-exon overexpression, suggesting that the broader transcript context is required for function in this case.

Taken together, these experiments confirm the fitness-promoting and fitness-neutral effects of exons within lncRNAs and demonstrate that the observed activity operates through the mature RNA transcript.

### Functional elements within NEAT1 and MALAT1 revealed by tiled pgRNAs

The dual CRISPR deletion strategy also enables high-resolution functional scanning of longer exons or unspliced transcripts. One well-studied example is Nuclear Enriched Abundant Transcript 1 (NEAT1), an unspliced lncRNA with two major isoforms: NEAT1_1 (short) and NEAT1_2 (long). NEAT1_2 serves as the architectural scaffold of paraspeckles, membraneless nuclear bodies involved in regulating gene expression and stress responses^31^. A previous study used CRISPR deletion to map NEAT1 functional domains, particularly for isoform switching (referred here as isoSwitch1 and isoSwitch2) and paraspeckle assembly. However, this study had limited resolution of 24 deletions^2^.

We performed a high-resolution functional scan of NEAT1 using 102 pgRNAs, with base-level coverage of ≥5 pgRNAs (Fig 5A). Most of the NEAT1 transcript is required for cell fitness, as indicated by negative log fold-change (LFC) of pgRNAs corresponding to deletions across the locus. Sequence-fitness mappings were significantly correlated between cell lines (Fig S4A). Our findings were broadly consistent with the earlier study: 1. pgRNAs overlapping isoSwitch2 showed higher impact on cell fitness than that of isoSwitch1 (Fig 5A); 2. deletion of the middle domain—previously shown to be necessary and sufficient for paraspeckle formation— led to decreased cell fitness. Notably, we also identified novel regions of pronounced fitness impact (arrows in the LFC plot Fig. 5A), raising the possibility of functional domains acting through paraspeckle-independent mechanisms.

**Figure 5:**
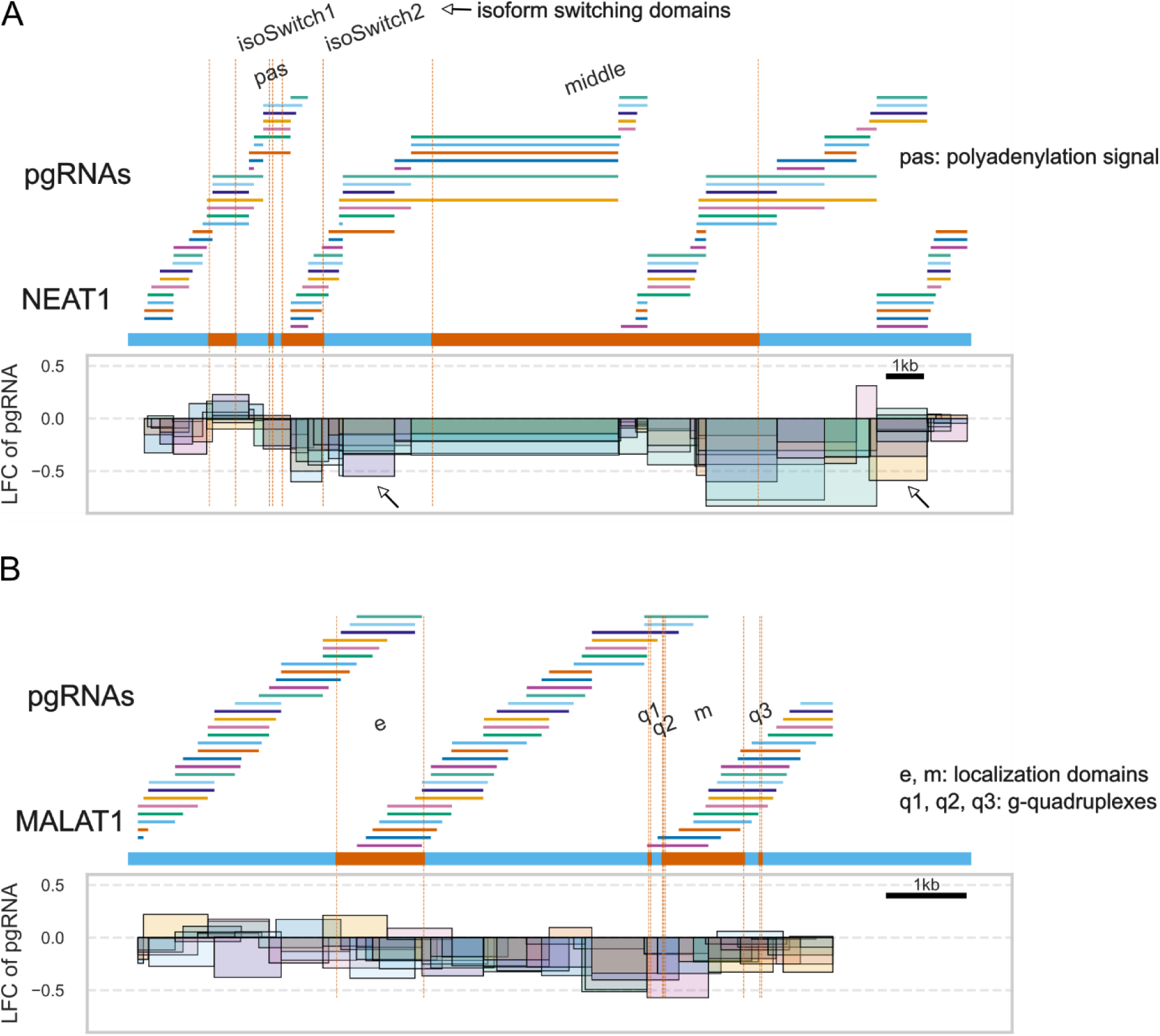
Functional elements in NEAT1 and MALAT1. (A) Illustration of pgRNAs tiled across NEAT1 gene (blue). Known functional domains are highlighted in orange. LFC of the pgRNAs is plotted below, each pgRNA is represented as a rectangle, height of which corresponds to the LFC while the width is proportional to the length. The relative position of the pgRNAs with respect to NEAT1 is maintained. Arrows represent pgRNAs with the lowest LFC values that do not overlap with the known NEAT1 domains. (B) Illustration of pgRNAs tiled across MALAT1 gene (blue). Known functional domains are highlighted in orange. LFC of the pgRNAs is plotted below.

We extended this approach to metastasis-associated lung adenocarcinoma transcript (MALAT1) (Fig 5B), abundant in the nuclear speckles. MALAT1 has two sequence elements (e and m), directing it to the nuclear speckles^32^. Three conserved RNA g-quadruplexes (q1, q2 and q3) have also been reported in the 3′ region, which are bound by Nucleolin and Nucleophosmin proteins and facilitate their localization to the nuclear speckles^33^. Interestingly, pgRNAs overlapping with these known domains had the lowest LFC in HeLa, implicating their role in cell fitness. The LFC of the pgRNAs (total 78) was correlated between the cell lines (Fig S4B), although to a lower extent than that of NEAT1. Overall, the high-resolution paired CRISPR method can be applied to map functional elements in longer transcripts at high resolution.

### Distribution and properties of lncRNA exons

We next leveraged these high-resolution maps to examine how functionality is distributed within lncRNA transcripts. To date, detailed organisational maps have been available for only a handful of lncRNAs, which generally suggest that functionality is confined to relatively narrow regions within the broader lncRNA transcript, encoding protein-binding sites or structured RNA domains^34,35^.

In our dataset, a small proportion of exons scored as hits in any given cell line, ranging from 3% to 22% (Fig S5A), corresponding to 1-2 exons per transcript (Fig 6A, S5B). In terms of total sequence, this corresponds to a global mean of approximately one-fifth of transcript nucleotides (Fig 6B), a figure that represents a lower bound, given the narrow phenotypic readout and false negative rate of the screen. Fitness-promoting exons were preferentially located toward the start of the transcript for all cell types (Fig 6C, S5C).

**Figure 6:**
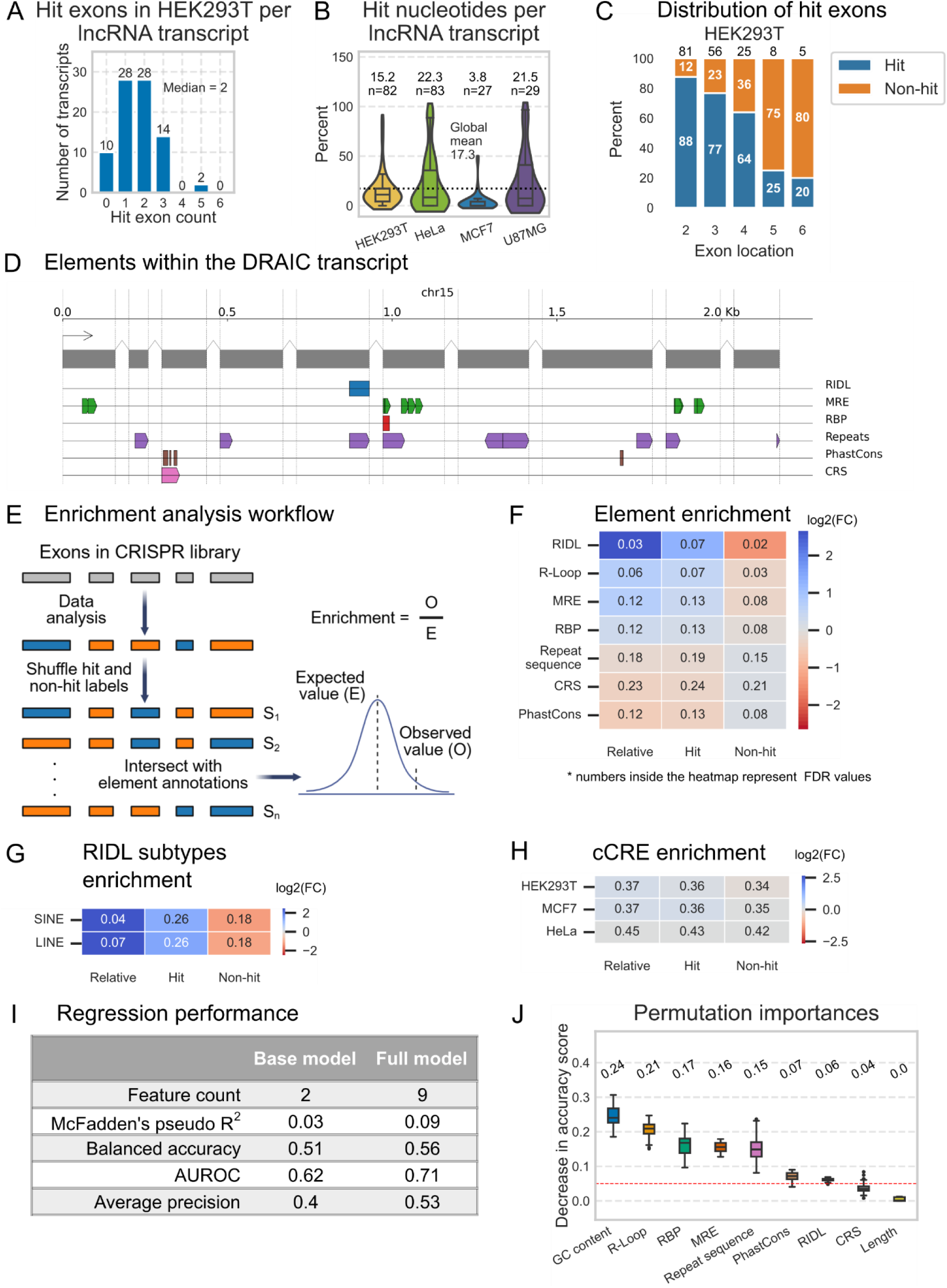
Properties of lncRNA exons. (A) Distribution of hit exon counts per transcript in hit lncRNAs. (B) Distribution of hit nucleotides per transcript in hit lncRNAs, the mean value is shown for each cell line and the number represents the sample size (total number of hit lncRNA transcripts). The global mean across all the cell lines is indicated by the dashed line. (C) Distribution of hit exons across transcript in hit lncRNAs, the numbers above the bars represent the total number of exons targeted at each location, aggregated from the entire transcript population. (D) Sequence elements in DRAIC exons (grey rectangles). Direction of the element indicates the strand information. (E) Workflow of enrichment analysis. The statistical significance and enrichment value were estimated by shuffling the hit and non-hit exon labels for 10,000 simulations, refer to the corresponding methods section for more details. (F) Heatmap colored by the log2 enrichment of sequence elements. Elements are indicated on the rows, the numbers inside the cells correspond to the FDR values. (G) Heatmap colored by the log2 enrichment of RIDL subtypes indicated on the rows (H)) Heatmap colored by the log2 enrichment values of cCREs for HEK293T, HeLa and MCF7 (cCRE annotations were not available for U87MG). (I) Comparison of logistic regression model trained on different sets of features. Base model was trained using GC content and length as features while full model uses overlap with the elements as additional features. (J) Importance of features for the trained model, estimated as a decrease in accuracy score. The red dashed line represents the 0.05 threshold.

Similar exon-mapping studies of protein coding genes have reported higher splicing inclusion levels for alternative fitness-promoting exons^36^, prompting us to investigate whether a similar effect governs lncRNAs. Surprisingly, this was observed only in HeLa cells (Fig S5D) and HeLa-specific hit exons, which are non-hits in other cell lines, did exhibit higher splicing inclusion in HeLa than in other cell lines (Fig S5E). Overall, the relatively lower expression of lncRNAs limits our ability to accurately quantify splicing inclusion.

We next asked whether exon functionality can be explained by sequence and structural elements^35^. Towards this, we curated the first comprehensive database of putative functional elements in lncRNAs, ElementaLdb (see Methods), incorporating both bioinformatically-predicted elements like repeat insertion domains of lncRNAs (RIDLs)^18,37^ and microRNA response elements (MREs)^38,39^ as well as experimentally-mapped elements like DNA:RNA hybrids (R-loops)^40^ and eCLIP-derived RBP interaction sites^41^. In total, ElementaLdb catalogues 18 million individual elements divided into 9 element types, 250 subtypes and is accessible at https://elementaldb.crg.eu/, where users can query, download and visualise lncRNA elements (Fig 6D).

To investigate the contribution of elements to exon functionality, we compared their per-nucleotide enrichment between hit and non-hit exons (Fig 6E). This revealed several significantly enriched element classes, including: RIDLs, R-loops, MREs and RBP sites (Fig 6F). RIDLs, a subset of repetitive elements^37^, showed the strongest enrichment, whereas repetitive sequences overall were depleted. Nine hit exons (8%) contained at least one RIDL, accounting for 2.7% of their total sequence. Fine-grained analysis revealed that 117 of 150 individual RBPs were significantly enriched in hits, and that SINE repeats were the most enriched RIDL subtype (Fig 6G). Element enrichment was consistent at the level of individual cell lines (Fig S5F).

Although the empirical enrichment test did not show significant enrichment of PhastCons conservation scores in the exons, splice sites of hit exons had higher PhastCons score than non-hits (Fig S5G), corroborating previous reports that evolutionary constraint operates on lncRNA splicing architecture^42^. Hit exons also displayed significantly higher GC content, while length distributions were comparable (Fig S5H).

As mentioned before, one possible confounder in our screen is the disruption of underlying regulatory DNA rather than the lncRNA transcript itself. To computationally evaluate this, we calculated the overlap between targeted exonic regions and DNA-encoded candidate cis-regulatory elements (cCREs)^43^. Using cell line-matched cCRE annotations from ENCODE, we observed no enrichment for cCREs amongst hit exons (Fig 6H), arguing against a contribution of DNA-encoded regulatory elements to the screen signal. Combined with the overexpression experiments (Fig 4F), these results support an RNA-mediated mechanism for observed hit exons.

We next investigated if the element annotations could classify exon functionality. A logistic regression model was trained using element overlap as features, with GC content and length as baseline covariates. Inclusion of element features provided a statistically significant but modest improvement over the baseline model (Fig 6I), yielding an AUROC of 0.7, balanced accuracy of 0.56 and average precision of 0.53. This demonstrates that sequence elements contribute meaningfully to exon functionality beyond the baseline predictive power of GC content and length alone.

GC content was the strongest individual predictor (Fig 6J), similar to fitness-promoting exons in protein coding genes^36^. While GC richness is a hallmark of lncRNA exons compared to introns^44^, higher GC content in fitness-promoting exons likely reflects a requirement for thermodynamic stability or secondary structures necessary for organization of the lncRNA. Other important features include DNA interactions (R-loops), RNA interactions (MREs) and protein interactions (RBP sites), supporting a role for these interactions in mediating lncRNA functionality.

### Sequence elements within fitness-promoting exons directly mediate lncRNA activity

To establish a direct causal link between specific elements and lncRNA function, we designed an experimental strategy based on overexpression of wild-type and mutant lncRNAs. To control for the potential confounding effects of altered transcript length and GC content arising from sequence deletion, we instead scrambled element sequences within an intact wild-type lncRNA background (Fig 7A, top). Three candidates were selected based on overlap with enriched element classes: (a) RIDL (SINE repeat) within DRAIC Exon 5, (b) an ILF3 (RBP) binding motif within LINC01270 Exon 2, (c) a U2AF2 (RBP) binding region and a PhastCons conserved sequence in LAMTOR5-AS1 Exon 2 (Fig 7A, bottom). As expected, WT transcripts promoted proliferation of HeLa cells relative to the empty vector control (Fig 7B). Scrambling of the target element completely abolished this fitness advantage in all three cases. Similar results were obtained in HEK293T cells (Fig 7C). This demonstrates that defined sequence elements within fitness-promoting exons are necessary for lncRNA bioactivity. It also highlights the potential of ElementaLdb combined with mutational analysis to identify lncRNA mechanisms, and add to existing evidence for the dependence of lncRNA function on surprisingly narrow sequence motifs^45^.

**Figure 7:**
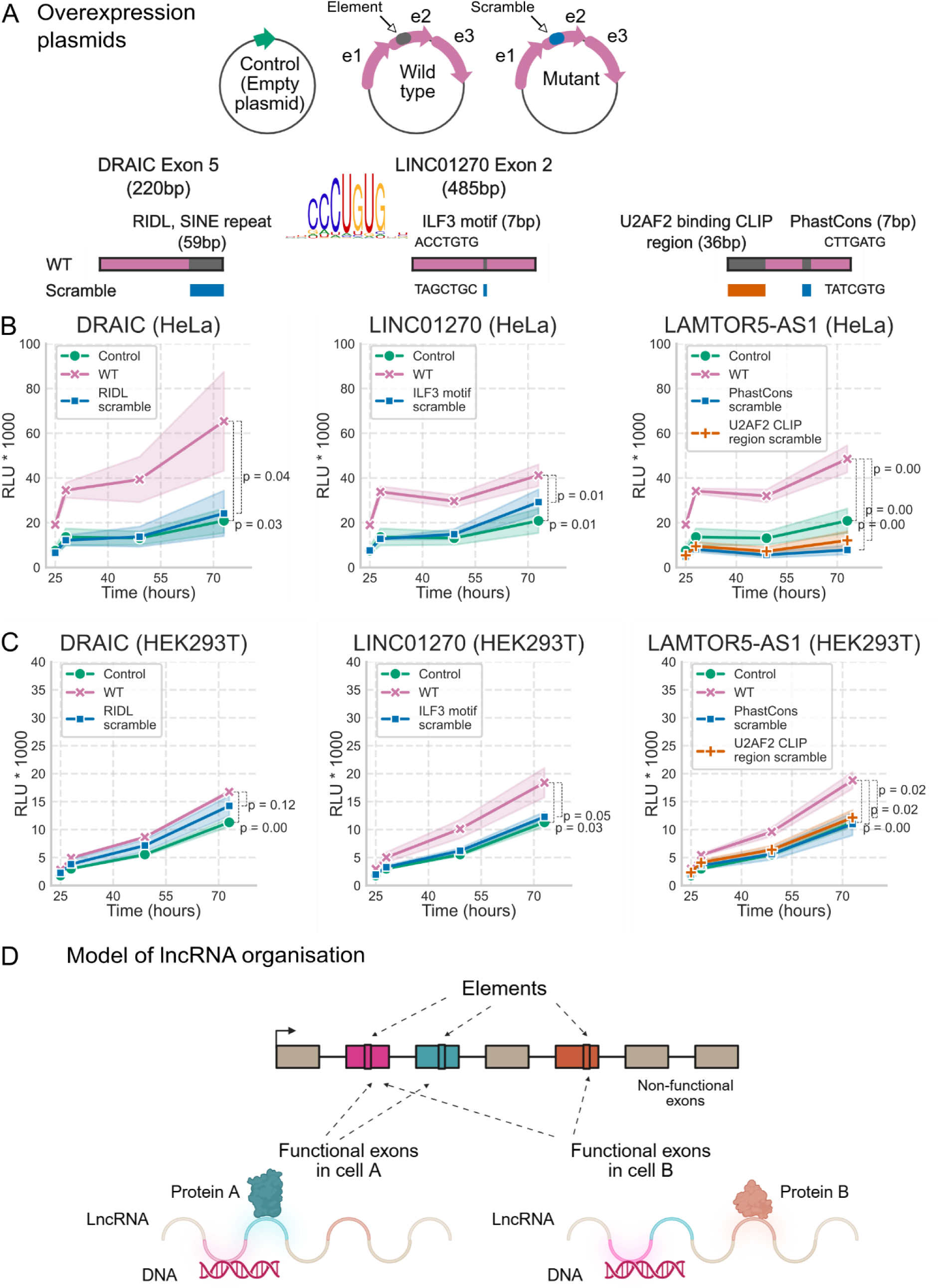
Elements within exons mediate lncRNA functionality. (A) Top: Design of overexpression plasmids. Bottom: schematic representation of the hit exon (pink) along with the sequence element highlighted in grey. The scrambled sequence is represented below the element. Growth curve of HeLa (B) and HEK293T(C) transfected with plasmids expressing the wild-type element, the scramble or the control. Proliferation was measured using CellTiterGlo luminescence values at 25h, 28h, 49h and 73h post transfection. Error bars indicate standard error with 3 biological replicates. Statistical significance was estimated using one-sided Student’s t-test by comparing area under the growth curve. (D) The proposed model of lncRNA organization. An example lncRNA with 7 exons is shown along with the sequence elements. Exons 2 and 3 are functional in cell type A, while exons 2 and 5 are functional in cell type B.

## Discussion

We present the most comprehensive experimental map of functional sequence in over 100 lncRNAs at exon-level resolution across four human cell lines. This is made possible by a high-throughput dual CRISPR-Cas9 deletion strategy, refined with inducible Cas9 and unique molecular identifiers. This pooled screening approach represents a dramatic improvement over state-of-the-art methods based on deletion mutant overexpression or individual CRISPR deletion, in terms of scale, resolution and hands-on time. Several independent lines of evidence supported the accuracy of this approach. Positive and negative controls indicated high sensitivity (Fig 2A, S2C) and specificity (Fig S2E), while pgRNAs targeting lncRNA TSSs served as internal positive controls that further supported screen performance (Fig 2C). The results also corroborated previously identified functional lncRNA exons, including those in PVT1 and HOTAIR.

The dropout screen design is inherently asymmetric in its sensitivity: while fitness-promoting exons are detected by progressive depletion of their targeting pgRNAs, exons whose deletion enhances fitness would manifest as pgRNA enrichment; a signal that is harder to distinguish from neutral noise in pooled screens^46^. Consequently, the “non-hit” category likely conflates genuinely dispensable exons with those that actively restrain cell fitness. Consistent with this, individual validation of several non-hit exons by competition assay revealed fitness-inhibiting effects upon deletion.

The functional maps deepen our understanding of lncRNA organisation. Fitness-promoting exons are not uniformly distributed within the transcripts; instead, they are preferentially located towards the 5′ end, mirroring the distribution of evolutionarily conserved patches reported previously^19^, while the majority of the transcript is dispensable for cell fitness. These observations are consistent with a modular lncRNA architecture, supporting the ‘beads-on-a-string’ model.

Our large dataset of hit and non-hit exons allowed us to compare their sequence properties and gain insights into underlying molecular mechanisms. Element scrambling experiments provided direct causal evidence linking specific sequence features to lncRNA bioactivity. Destruction of the enriched sequence elements in the three tested exons abolished lncRNA activity. The exon-resolution functional maps, together with the ElementaLdb database, thus provide a rich resource for generating testable hypotheses regarding lncRNA mechanisms of action.

Our findings point to diverse molecular mechanisms for lncRNA elements that broadly support previous reports: (1) The SINE family of transposable elements contribute to lncRNA functionality through roles in subcellular localization^17^ and translation regulation of a bound mRNA^47^, although the precise mechanism of the MIR SINE element in DRAIC Exon 5 requires further investigation. (2) The loss of LINC01270 activity upon scrambling of the ILF3 motif establishes a previously unreported functional interaction between this lncRNA and ILF3. ILF3 is associated with poor prognosis^48^ in colorectal cancer, suggesting that the fitness role of LINC01270 may be mediated through ILF3. (3) Similarly, LAMTOR5-AS1 function depended on a U2AF2 binding region. U2AF2 is a component of the heterodimeric splicing factor U2AF^49^, which also has splicing-independent roles like RNA export and translational regulation^50^. This raises the possibility that U2AF2 binding regions within lncRNAs reflect the formation of functional ribonucleoprotein complexes rather than mere RNA processing signals.

Integrating our observations, we propose a model of lncRNA organisation in which transcripts comprise a mixture of functional and dispensable exons (Fig 7D). The functionality of individual exons is governed by the sequence elements, and which exons are functionally engaged depends on the proteomic landscape of the cell. This context-dependent modularity offers a parsimonious explanation for the frequently conflicting reports of lncRNA function across different cell types, and underscores the necessity of sub-genic mapping: a mutation in one exon may be pathogenic in a specific cellular context while remaining silent in others.

Despite these significant advancements, our study highlights several areas for further investigation. (1) While the screen focused on cell fitness, other phenotypic outputs, such as differentiation or migration, may reveal additional functional domains. The higher conservation scores of splice sites of non-hit exons (Fig S5G) indeed points to a population of exons which are either incorrectly called as non-hits or have functions beyond cell fitness. (2) Future iterations utilizing RNA-targeting CRISPR systems such as Cas13^51^ will be valuable for decoupling the contributions of the DNA locus from the RNA transcript. (3) Finally, while the element-based model captures a significant fraction of the variance in exon function, it explains only part of the functionality, pointing to incomplete annotations or additional layers of regulation that remain to be systematically explored.

In conclusion, the exon-resolution functional maps substantially expand the catalogue of experimentally characterised lncRNA functional domains. Along with the ElementaLdb database, these functional maps provide a framework for prioritising non-coding variants in disease-associated loci and for the rational design of RNA-based therapeutics.

## Materials & Methods

### Cell culture

HEK293T (female), MCF7 (female), and HeLa (female) cell lines were generously provided by Adrian Ochsenbein’s group at the University Hospital of Bern, Switzerland. The U87MG cell line with male epithelial morphology was obtained from the American Type Culture Collection (ATCC HTB-14). Authentication of all cell lines was performed using Short Tandem Repeat (STR) profiling (Microsynth Cell Line Typing), and they were tested negative for mycoplasma contamination. HEK293T, HeLa, and MCF7 cells were cultured in DMEM, while U87MG cells were cultured in EMEM. All media were supplemented with 10% Fetal Bovine Serum, 1% L-Glutamine, and 1% Penicillin-Streptomycin. Cells were split every 2–3 days and maintained at 37°C in a humid atmosphere with 5% CO_2_.

### Lentivirus production

Lentivirus were produced by co-transfecting HEK293T cells with the following plasmids: 12.5 μg of DD-Cas9 plasmid with zeocin resistance (Addgene #90085) lacking the promoter and downstream guide RNA, 7.5 μg of psPAX2 plasmid, and 4 μg of the packaging pVsVg plasmids. The transfection was performed using Lipofectamine2000 (ThermoFisher #11668019). Prior to transfection, 2.5 million HEK293T cells were seeded in a 10 cm dish coated with Poly-L-Lysine (Sigma #P4832) diluted 1:5 in 1X PBS. The supernatant containing viral particles was harvested at 24, 48, and 72 hours post-transfection. Viral particles were concentrated 100-fold by adding 1 volume of cold PEG-it Virus Precipitation Solution (BioCat #LV810A-1-SBI) to every four volumes of supernatant. After incubation at 4°C for 12 hours, the supernatant/PEG-it mixture was centrifuged at 1,500 × g for 30 minutes at 4°C. The resulting pellet was resuspended in 1X PBS, stored at −80°C, and used as needed.

### Stable cell line production

To generate stable DD-Cas9-expressing cell lines, HeLa cells were incubated for 24 hours with a culture medium containing a concentrated viral preparation carrying pLentiDDCas9-P2A-mVenus and 8 μg/mL polybrene. Infected cells were selected for at least five days with zeocin (70 μg/mL) and then subjected to fluorescence-activated cell sorting (FACS) to obtain a population with at least 60% mVenus-positive cells.

HEK293T, MCF7, and U87MG cell lines were co-transfected with a PiggyBac transposon vector containing DD-Cas9, BFP, and blasticidin resistance (designed in-house and synthesized by VectorBuilder) and the PiggyBac transposase pCMV-hyPBase (a gift from Sanger Institute) at a 10:1 ratio of transposon to transposase. Transfection was carried out using Lipofectamine2000 (ThermoFisher #11668019) for HEK293T and U87MG cells, and Lipofectamine 3000 (ThermoFisher #L3000001) for MCF7 cells. Transfected cells were selected for at least five days with blasticidin (5 μg/mL) and then subjected to FACS sorting to obtain a population with at least 60% BFP-positive cells.

### Lentiviral titer calculation and infection

To achieve the desired multiplicity of infection (MOI) of 1, a titration experiment was performed in HeLa, HEK293T, U87MG, and MCF7 cell lines. A 12-well plate was seeded with 2 million cells per well and supplemented with 8 μg/mL polybrene. The virus containing the library was added to each well in volumes ranging from 0.5 to 2 μL and transduction was performed using spin-infection (centrifuging at 1500 rpm and 37°C for 90 minutes). After centrifugation, the media was replaced with fresh media without polybrene and incubated overnight. The following day, cells were counted, and each well was split into two equal aliquots, of which one was treated with 2 μg/mL puromycin. After 72 hours, the MOI was calculated by dividing the number of surviving cells in the puromycin-treated well by the number in the puromycin-free well. An MOI of 1 was used for all screening experiments. For large-scale screens, 20 million cells were seeded in 12-well plates at a density of 2 million cells per well for spin-infection. The following day, cells were pooled together, and fresh media containing puromycin (2 μg/mL) was added. Puromycin selection was maintained for six days until phenotypic screens were initiated.

### CRISPR-deletion efficiency

DD-Cas9-expressing HeLa, HEK293T, MCF7 and U87MG were transduced with a pDECKO^52^ plasmid containing targeting and non-targeting pgRNAs as described above (Lentiviral titer calculation and infection). Non-targeting pgRNAs are directed against the *GFP* gene and targeting pgRNAs are instead aimed at the *MALAT1* gene enhancer. The following day, each condition was split into two 10 cm dishes, both selected with 2 μg/mL puromycin, and one of which was treated with 400 nM Shield-1. Cells were incubated at 37°C in a humidified, 5% CO_2_ atmosphere for 72h, after which they were harvested and their genomic DNA was extracted.

qPCRBIO SyGreen 1-Step Detect Lo-ROX kit (PCRBIOSYSTEMS #PB25.11-01) was used to calculate the non-deleted alleles ratio for each condition and cell line. The reaction was conducted in duplicates with one primer inside and another one outside the deletion. CT values of the target gene (*MALAT1*) were normalized to the CT values of the housekeeping control (*GAPDH*) using the ΔΔCT method.

### Genomic DNA cleavage

Cas9 expressing HeLa cells were transfected using Lipofectamine2000 (ThermoFisher #11668019) or transduced with DECKO plasmids carrying the indicated pgRNAs. Cells were selected for 2 days with 2 μg/mL puromycin, after which cells were pelleted, and genomic DNA was extracted using the Thermo Scientific™ GeneJET Genomic DNA Purification Kit (ThermoFisher #K0721) following the manufacturer’s protocol. PCR was performed with genomic primers flanking the deleted region using the DreamTaq Green PCR Master Mix (2X) (ThermoFisher #K1081). Touchdown PCR cycling conditions were applied. The PCR products were analyzed on agarose gel (% depending on amplicons size).

### Candidate selection and library design

We generated a full annotation file by merging Gencode v38 and MiTranscriptome v2. MiTranscriptome file was lifted over from hg19 to hg38 with an in-house script. In short, a BED file with all exons in the annotation file was lifted over with LiftOver. Then, each individual transcript ID was checked for correct structure, meaning that transcripts with any of these coordinate transformations were removed: inversion, splitting, unmapped, and exons rearranged compared to those in the original transcript. Exons were discarded if they overlapped with a protein coding exon according to Gencode annotation. We prioritized lncRNAs known be required for cell fitness, leading to a total of 107 lncRNAs, 29 from Cancer LncRNA Census 2^26^ and 78 from previous CRISPRi screen^22^. To avoid redundancy in targeted regions, we combined groups of exons targeting the same region. We used CRISPETa^53^ to find pgRNAs targeting these regions. This designed lentiviral library was then obtained pre-packaged from VectorBuilder. Validated pgRNAs were cloned into DECKO plasmids in-house.

### CRISPR screen

One week after infection (time point 0 or T_0_), cells were counted, and the reference sample (t_0_, 12 million cells, corresponding to a library coverage of approximately 1,000x) was collected. Cells were cultured in 150 mm culture-treated dishes, and 12 million cells were passed every 2-3 days to maintain a coverage of approximately 1,000X (defined as the number of cells divided by the number of unique library sequences). At T_0_, 400 nM Shield-1 (MedChemExpress #HY-112210) was added to the cell culture for 3 days to stabilize the DD-Cas9 protein, and 300 nM M3814 (MedChemExpress #HY-101570) was added for 18 hours to enhance CRISPR deletion. Cells were harvested at 7 (T_1_), 14 (T_2_), and 21 (T_3_) days following T_0_ for genomic DNA extraction.

### Screen analysis and identification of hit exons

MAGeCK^54^ was used to identify hit regions (FDR <= 0.1) at each time point with half of the negative controls (38) used for normalization. The regions were mapped to overlapping exons of lncRNAs using BEDTools^55^. An exon was considered to be hit if at least one overlapping region was hit. Exons of transcripts with expression more than 0.1 TPM in all the replicates were considered so that only consistently expressed regions are retained. If a gene did not have any transcript satisfying this criterion, the longest transcript of the gene was used if the gene TPM passed the same filtering condition. For exon level analysis, exons that are at least 20% overlapping with each other were merged using BEDTools and long exons (length above 2000bp) were filtered out.

### Genomic DNA preparation and sequencing

Genomic DNA (gDNA) was isolated using the Blood & Cell Culture DNA Midi kit (Qiagen #13343) following the manufacturer’s instructions. The concentration of gDNA was determined using a Nanodrop. For PCR amplification, gDNA was divided into 50 μL reactions, ensuring that each well contained at most 2 μg of gDNA. PCR was performed using NEBNext® Ultra™ II Q5® Master Mix (NEB #M0544), universal forward primer and a uniquely barcoded P7 reverse primer. PCR cycling conditions were as follows: initial denaturation at 98°C for 30 seconds, followed by 25 cycles of denaturation at 98°C for 10 seconds, annealing at 52°C for 30 seconds, extension at 72°C for 30 seconds, and a final extension at 72°C for 2 minutes. PCR products were purified using SPRIselect beads at a ratio of 0.625 to the sample, following the manufacturer’s instructions (Beckman Coulter #B23318). The purified PCR products were quantified using the Qubit dsDNA HS Assay Kit (ThermoFisher #Q32854). The quality of the samples was assessed using the Bioanalyzer and Agilent High Sensitivity DNA Kit (Agilent #5067-4626). The samples were sequenced on a HiSeq2000 (Illumina) with paired-end 150 bp reads, targeting a coverage of 40 million reads per sample.

### Post-cleavage RNA splicing

Cas9 expressing HeLa cells were transfected with DECKO plasmids carrying the indicated pgRNAs using Lipofectamine2000 (ThermoFisher #11668019). Cells were selected for 2 days with 2 μg/mL puromycin, after which cells were pelleted, and total RNA was extracted using the Thermo Scientific™ GeneJET RNA Purification Kit (ThermoFisher #K0731) following the manufacturer’s protocol. The RNAs were reverse transcribed to produce cDNAs using specific primers and the GoTaq® 1-Step RT-qPCR kit (Promega #A6020), following the manufacturer’s protocol. The cDNAs were then used for nested PCR with the DreamTaq Green PCR Master Mix (2X) (ThermoFisher #K1081) and indicated primers. The PCR products were analyzed on agarose gel (% depending on amplicons size). The bands were extracted from the agarose gels using the GeneJET Gel Extraction and DNA Cleanup Micro Kit (ThermoFisher #K0832) following the manufacturer’s protocol. The extracted bands were sent for Sanger sequencing to Eurofins.

### Competition assay

HeLa cells were infected with DECKO lentivirus expressing fluorescent proteins. The virus expressing control pgRNAs targeting AAVS1 expressed GFP (pgRNAs-GFP+), while the pgRNAs targeting candidate regions expressed mCherry (pgRNAs-mCherry+). After infection and two days of puromycin selection (2 μg/mL), GFP and mCherry cells were mixed 1:1 in a 6-well plate (total of 100,000 cells). Cell counts were analyzed by CytoFLEX LX (Beckman Coulter) on day 0, 3, 5 and 9, when cells were split on a 1:3 ratio.

### RNA sequencing analysis

Public RNAseq data was downloaded from SRA using these accessions numbers: HEK293T: SRR1573494, SRR1573495; HeLa: SRR24928373, SRR24928374, SRR24928375; MCF7: SRR32118610, SRR32118611, SRR32118612; U87MG: SRR12282424, SRR12282426, SRR12282428. Analysis was done using nf-core/rnaseq pipeline (version 3.18.0)

### Overexpression proliferation assay

Cells per well were seeded in a 96-Well Clear Bottom White Polystyrene Microplate (Corning #10517742) at 20% confluency (1.5k cells). Sixteen hours post-plating, cells were transfected in triplicates with 50 ng of pcDNA3.1 expressing the indicated gene using 0.2 µL of Lipofectamine2000 (ThermoFisher #11668019) per well for HeLa, HEK293T and U87MG, and 0.2 µL of Lipofectamine3000 (ThermoFisher #L3000015) for MCF7. The media was replaced after 6 hours of transfection.

Proliferation was measured using RealTime-Glo MT Cell Viability Assay (Promega #G9712) according to the manufacturer’s protocol. Twenty-four hours after transfection, the media was changed to complete media supplemented with 1x MT Cell Viability Substrate and 1x NanoLuc® Enzyme. After incubating the cells for 1h at 37°C in a humidified, 5% CO_2_ atmosphere, the luminescence was measured at 37 °C with Tecan Infinite 200 Pro at different time points.

### Tiling plot

The tiling guides of NEAT1 and MALAT1 correspond to the regions R1111 and R1110 respectively. The LFC at the last time point was used in the plots.

### Exon splicing quantification

SpliSER^56^ was used to calculate the Splice-site Strength Estimates (SSE) for all the splice sites (exon junctions). Splice sites with less than 5 coverage were filtered out, since SSE is not reliable in such cases. SSE values across replicates were aggregated as median values. Mean of all the SSE of the splice sites overlapping with the merged exon was used to compare the splicing levels.

### Enrichment analysis

The following putative lncRNA sequence elements were obtained from the ElementaLdb server: Conserved RNA structures (CRS)^57^, microRNA response elements (MRE), PhastCons^58^, RBP sites, repeat sequences, RIDL^18^, R-Loop. To calculate enrichment of these annotations in the exons, the expected overlap was estimated by shuffling the hit and non-hit labels, an approach similar to the Genomic Association Test^59^. The exon labels were shuffled while maintaining the total length within 0.1% range of the observed overlap. The median of percent overlaps of 10000 shuffles was used as the expected overlap and significance was estimated based on the FDR corrected q values from the distribution. To calculate relative enrichment in hits compared to non-hits, the distribution of ratio of overlap of hit and non-hits for each simulation was used instead of the percent overlap. For aggregate level enrichment analysis, combined hits from all the four cell lines were used. To further analyse cell line specific hits, we obtained chromatin associated RNA data for HEK293T using SRA accession number SRR8206679.

### Conservation of splice sites

Average PhastCons (470way, Michael Hiller lab) score in 5bp window around the exon intron boundary were computed separately for donor and acceptor sites. Mean of all the splice sites scores overlapping with the merged exon was used to compare the splice site conservation. Matching dinucleotides farthest within a 200-500 bp region away from the exon boundary were used as controls.

### Linear regression

The percent overlap of exons was calculated with the above annotations. A logistic regression model was trained using scikit-learn package, using these overlap values along with GC content and length as features. The performance of this full model was compared with the base model with only GC content and length as features. The significance was estimated using likelihood ratio test, comparing the likelihood of full model with the base model. Feature importance was calculated using sklearn.inspection.permutation_importance.

## Acknowledgements

The authors thank the members of GOLD Lab and Marsico Lab for insightful feedback and discussions. The authors acknowledge administrative and logistical support from Basak Ginsbourger, Rahel Tschudi, Marla Rittiner, Beatrice Stalder, Willy Hofstetter and Patrick Furer (DBMR, University of Bern). Computations were carried out on the University of Bern Interfaculty Bioinformatics Unit cluster maintained by Rémy Bruggman and Pierre Berthier, and UCD Sonic High Performance Computing (HPC) cluster maintained by UCD Research IT. This work was funded by the Swiss National Science Foundation through the National Centre of Competence in Research (NCCR) “RNA & Disease” (51NF40-182880), project funding “The elements of long noncoding RNA function” (31003A_182337), Sinergia project “Regenerative strategies for heart disease via targeting the long noncoding transcriptome” (173738); by the Medical Faculty of the University and University Hospital of Bern; by the Helmut Horten Stiftung, Swiss Cancer Research Foundation (4534-08-2018); and by Science Foundation Ireland through Future Research Leaders award 18/FRL/6194 (to R.J.). This research was also funded by Science Foundation Ireland under Grant number [18/CRT/6214] (to S.B.) and in part by the EU’s Horizon 2020 research and innovation programme under the Marie Sklodowska-Curie grant H2020-MSCA-COFUND-2019-945385.

## References

1. Mattick, J. S. et al. Long non-coding RNAs: definitions, functions, challenges and recommendations. Nat. Rev. Mol. Cell Biol. 24, 430–447 (2023).

2. Yamazaki, T. et al. Functional Domains of NEAT1 Architectural lncRNA Induce Paraspeckle Assembly through Phase Separation. Mol. Cell 70, 1038–1053.e7 (2018).

3. Cerase, A., Pintacuda, G., Tattermusch, A. & Avner, P. Xist localization and function: new insights from multiple levels. Genome Biol. 16, 166 (2015).

4. Guttman, M. & Rinn, J. L. Modular regulatory principles of large non-coding RNAs. Nature 482, 339–346 (2012).

5. Wutz, A., Rasmussen, T. P. & Jaenisch, R. Chromosomal silencing and localization are mediated by different domains of Xist RNA. Nat. Genet. 30, 167–174 (2002).

6. Li, Y. et al. A noncoding RNA modulator potentiates phenylalanine metabolism in mice. Science 373, 662–673 (2021).

7. Jin, F. et al. A functional motif of long noncoding RNA Nron against osteoporosis. Nat. Commun. 12, 3319 (2021).

8. Plaisance, I. et al. A transposable element into the human long noncoding RNA CARMEN is a switch for cardiac precursor cell specification. Cardiovasc. Res. 119, 1361–1376 (2023).

9. Farberov, S. et al. Structural features within the NORAD long noncoding RNA underlie efficient repression of Pumilio activity. Nat. Struct. Mol. Biol. 32, 287–299 (2025).

10. Uroda, T. et al. Conserved Pseudoknots in lncRNA MEG3 Are Essential for Stimulation of the p53 Pathway. Mol. Cell 75, 982–995.e9 (2019).

11. De Souza Degenhardt, M. F. & Wang, Y.-X. Challenges and opportunities in technologies and methods for lncRNA structure determination. Cell Biosci. 15, 132 (2025).

12. Van Nostrand, E. L. et al. Robust transcriptome-wide discovery of RNA-binding protein binding sites with enhanced CLIP (eCLIP). Nat. Methods 13, 508–514 (2016).

13. Tichon, A., Perry, R. B.-T., Stojic, L. & Ulitsky, I. SAM68 is required for regulation of Pumilio by the NORAD long noncoding RNA. Genes Dev. 32, 70–78 (2018).

14. Dehingia, B. et al. RNA-binding proteins mediate the maturation of chromatin topology during differentiation. Nat. Cell Biol. 27, 1510–1525 (2025).

15. Kallen, A. N. et al. The Imprinted H19 LncRNA Antagonizes Let-7 MicroRNAs. Mol. Cell 52, 101–112 (2013).

16. Niehrs, C. & Luke, B. Regulatory R-loops as facilitators of gene expression and genome stability. Nat. Rev. Mol. Cell Biol. 21, 167–178 (2020).

17. Lubelsky, Y. & Ulitsky, I. Sequences enriched in Alu repeats drive nuclear localization of long RNAs in human cells. Nature 555, 107–111 (2018).

18. Carlevaro-Fita, J. et al. Ancient exapted transposable elements promote nuclear enrichment of human long noncoding RNAs. Genome Res. 29, 208–222 (2019).

19. Hezroni, H. et al. Principles of Long Noncoding RNA Evolution Derived from Direct Comparison of Transcriptomes in 17 Species. Cell Rep. 11, 1110–1122 (2015).

20. Ross, C. J. et al. Uncovering deeply conserved motif combinations in rapidly evolving noncoding sequences. Genome Biol. 22, 29 (2021).

21. Tsai, M.-C. et al. Long Noncoding RNA as Modular Scaffold of Histone Modification Complexes. Science 329, 689–693 (2010).

22. Liu, S. J. et al. CRISPRi-based genome-scale identification of functional long noncoding RNA loci in human cells. Science 355, eaah7111 (2017).

23. Esposito, R. et al. Multi-hallmark long noncoding RNA maps reveal non-small cell lung cancer vulnerabilities. Cell Genomics 2, 100171 (2022).

24. Cho, S. W. et al. Analysis of off-target effects of CRISPR/Cas-derived RNA-guided endonucleases and nickases. Genome Res. 24, 132–141 (2014).

25. Senturk, S. et al. Rapid and tunable method to temporally control gene editing based on conditional Cas9 stabilization. Nat. Commun. 8, 14370 (2017).

26. Vancura, A. et al. Cancer LncRNA Census 2 (CLC2): an enhanced resource reveals clinical features of cancer lncRNAs. NAR Cancer 3, zcab013 (2021).

27. Yao, Q., Zhang, X. & Chen, D. The emerging potentials of lncRNA DRAIC in human cancers. Front. Oncol. 12, (2022).

28. Pal, G. et al. Long Noncoding RNA from PVT1 Exon 9 Is Overexpressed in Prostate Cancer and Induces Malignant Transformation and Castration Resistance in Prostate Epithelial Cells. Genes 10, 964 (2019).

29. Li, C. et al. Long noncoding RNA plasmacytoma variant translocation 1 is overexpressed in cutaneous squamous cell carcinoma and exon 2 is critical for its oncogenicity. Br. J. Dermatol. 190, 415–426 (2024).

30. Wu, L., Murat, P., Matak-Vinkovic, D., Murrell, A. & Balasubramanian, S. Binding Interactions between Long Noncoding RNA HOTAIR and PRC2 Proteins. Biochemistry 52, 9519–9527 (2013).

31. Smith, N. E., Spencer-Merris, P., Fox, A. H., Petersen, J. & Michael, M. Z. The Long and the Short of It: NEAT1 and Cancer Cell Metabolism. Cancers 14, 4388 (2022).

32. Miyagawa, R. et al. Identification of cis- and trans-acting factors involved in the localization of MALAT-1 noncoding RNA to nuclear speckles. RNA 18, 738–751 (2012).

33. Ghosh, A. et al. Identification of G-quadruplex structures in MALAT1 lncRNA that interact with nucleolin and nucleophosmin. Nucleic Acids Res. 51, 9415–9431 (2023).

34. Quinn, J. J. et al. Revealing long noncoding RNA architecture and functions using domain-specific chromatin isolation by RNA purification. Nat. Biotechnol. 32, 933–940 (2014).

35. Gandhi, M., Caudron-Herger, M. & Diederichs, S. RNA motifs and combinatorial prediction of interactions, stability and localization of noncoding RNAs. Nat. Struct. Mol. Biol. 25, 1070–1076 (2018).

36. Xiao, M.-S. et al. Genome-scale exon perturbation screens uncover exons critical for cell fitness. Mol. Cell 84, 2553–2572.e19 (2024).

37. Johnson, R. & Guigó, R. The RIDL hypothesis: transposable elements as functional domains of long noncoding RNAs. RNA 20, 959–976 (2014).

38. Jeggari, A., Marks, D. S. & Larsson, E. miRcode: a map of putative microRNA target sites in the long non-coding transcriptome. Bioinformatics 28, 2062–2063 (2012).

39. Li, J.-H., Liu, S., Zhou, H., Qu, L.-H. & Yang, J.-H. starBase v2.0: decoding miRNA-ceRNA, miRNA-ncRNA and protein–RNA interaction networks from large-scale CLIP-Seq data. Nucleic Acids Res. 42, D92–D97 (2014).

40. Sanz, L. A. et al. Prevalent, Dynamic, and Conserved R-Loop Structures Associate with Specific Epigenomic Signatures in Mammals. Mol. Cell 63, 167–178 (2016).

41. Van Nostrand, E. L. et al. A large-scale binding and functional map of human RNA-binding proteins. Nature 583, 711–719 (2020).

42. Nitsche, A., Rose, D., Fasold, M., Reiche, K. & Stadler, P. F. Comparison of splice sites reveals that long noncoding RNAs are evolutionarily well conserved. RNA 21, 801–812 (2015).

43. Moore, J. E. et al. An expanded registry of candidate cis-regulatory elements. Nature 1–10 (2026) doi:10.1038/s41586-025-09909-9.

44. Haerty, W. & Ponting, C. P. Unexpected selection to retain high GC content and splicing enhancers within exons of multiexonic lncRNA loci. RNA 21, 320–332 (2015).

45. Zhang, B. et al. A Novel RNA Motif Mediates the Strict Nuclear Localization of a Long Noncoding RNA. Mol. Cell. Biol. 34, 2318–2329 (2014).

46. Doench, J. G. Am I ready for CRISPR? A user’s guide to genetic screens. Nat. Rev. Genet. 19, 67–80 (2018).

47. Schein, A., Zucchelli, S., Kauppinen, S., Gustincich, S. & Carninci, P. Identification of antisense long noncoding RNAs that function as SINEUPs in human cells. Sci. Rep. 6, 33605 (2016).

48. Li, K. et al. ILF3 is a substrate of SPOP for regulating serine biosynthesis in colorectal cancer. Cell Res. 30, 163–178 (2020).

49. Glasser, E. et al. Pre-mRNA splicing factor U2AF2 recognizes distinct conformations of nucleotide variants at the center of the pre-mRNA splice site signal. Nucleic Acids Res. 50, 5299–5312 (2022).

50. Garcia, G. R. et al. U2AF regulates the translation and localization of nuclear-encoded mitochondrial mRNAs. 2024.09.18.613780 Preprint at 10.1101/2024.09.18.613780 (2024).

51. Sun, Q. et al. Systematic screening for functional exon-skipping isoforms using the CRISPR-RfxCas13d system. Cell Syst. 16, (2025).

52. Aparicio-Prat, E. et al. DECKO: Single-oligo, dual-CRISPR deletion of genomic elements including long non-coding RNAs. BMC Genomics 16, 846 (2015).

53. Pulido-Quetglas, C. et al. Scalable Design of Paired CRISPR Guide RNAs for Genomic Deletion. PLOS Comput. Biol. 13, e1005341 (2017).

54. Li, W. et al. MAGeCK enables robust identification of essential genes from genome-scale CRISPR/Cas9 knockout screens. Genome Biol. 15, 554 (2014).

55. Quinlan, A. R. BEDTools: The Swiss-Army Tool for Genome Feature Analysis. Curr. Protoc. Bioinforma. 47, 11.12.1–11.12.34 (2014).

56. Dent, C. I. et al. Quantifying splice-site usage: a simple yet powerful approach to analyze splicing. NAR Genomics Bioinforma. 3, lqab041 (2021).

57. Seemann, S. E. et al. The identification and functional annotation of RNA structures conserved in vertebrates. Genome Res. 27, 1371–1383 (2017).

58. Siepel, A. et al. Evolutionarily conserved elements in vertebrate, insect, worm, and yeast genomes. Genome Res. 15, 1034–1050 (2005).

59. Heger, A., Webber, C., Goodson, M., Ponting, C. P. & Lunter, G. GAT: a simulation framework for testing the association of genomic intervals. Bioinformatics 29, 2046–2048 (2013).

