## supplementary info for "Organisational principles of long non-coding RNAs revealed by exon deletion"

### Supplementary Figures

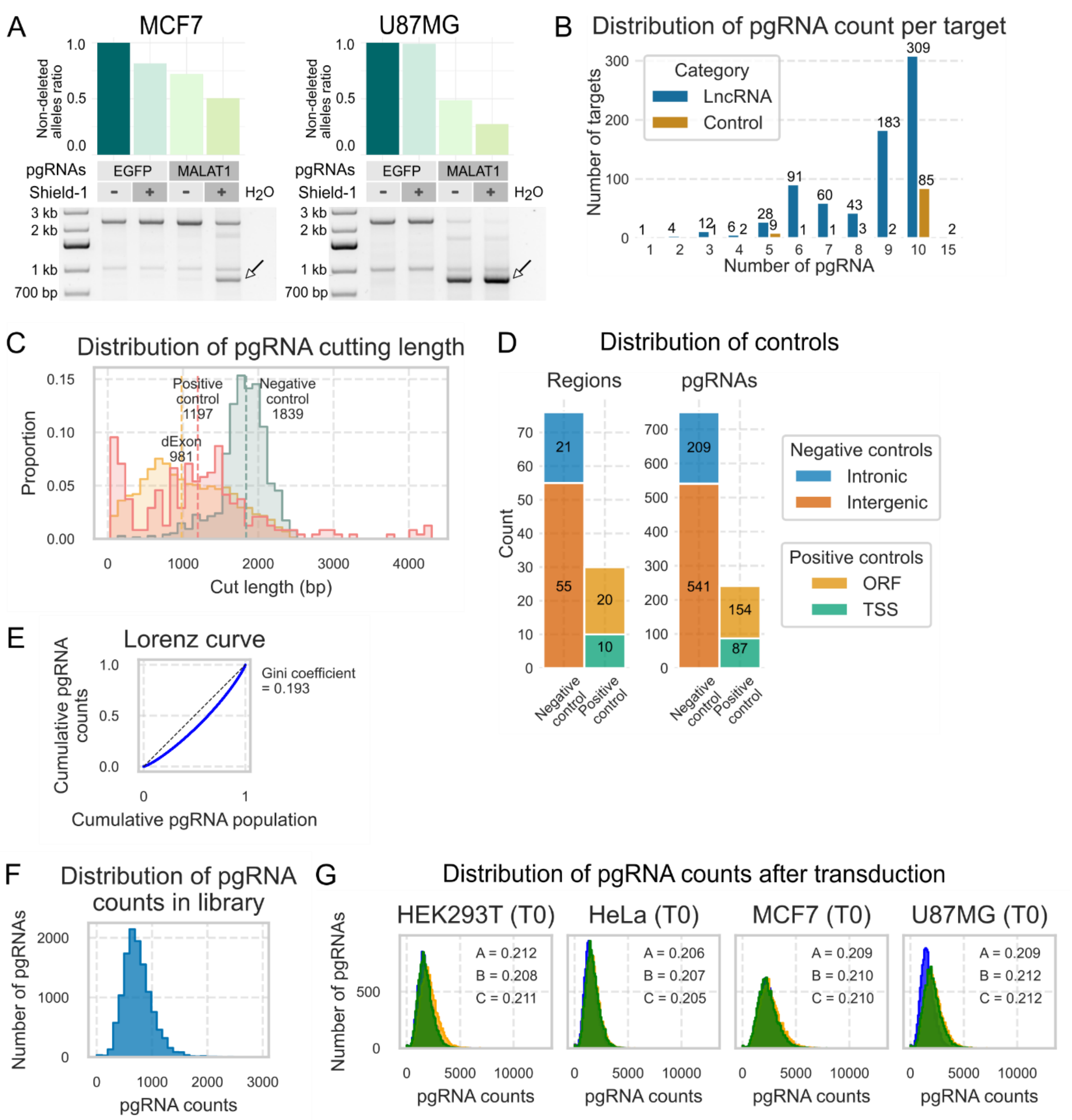


*Figure S1: High-resolution CRISPR screen to study lncRNA exons. (A) Top: Non-deleted alleles ratio corresponding to pgRNAs targeting the MALAT1 enhancer and non-targeting EGFP control, with and without Shield-1 treatment in MCF7 and U87MG cell lines. Bottom: Gel electrophoresis showing the wild type and genomic deletion amplicon of MALAT1 enhancer. The arrows indicate the DNA fragment with the expected deletion. (B) Distribution of the number of pgRNAs targeting each genomic region. (C) Distribution of the cutting length of the regions for positive controls, negative controls and lncRNA regions. (D) Number of target regions and corresponding pgRNAs for positive and negative controls. (E) Lorenz curve of the CRISPR library. (F) Distribution of pgRNA counts in the library. (G) Distribution of pgRNA counts at the initial timepoint after transduction for each cell line tested. For each replicate A, B and C, the gini coefficient is shown in the plot.*

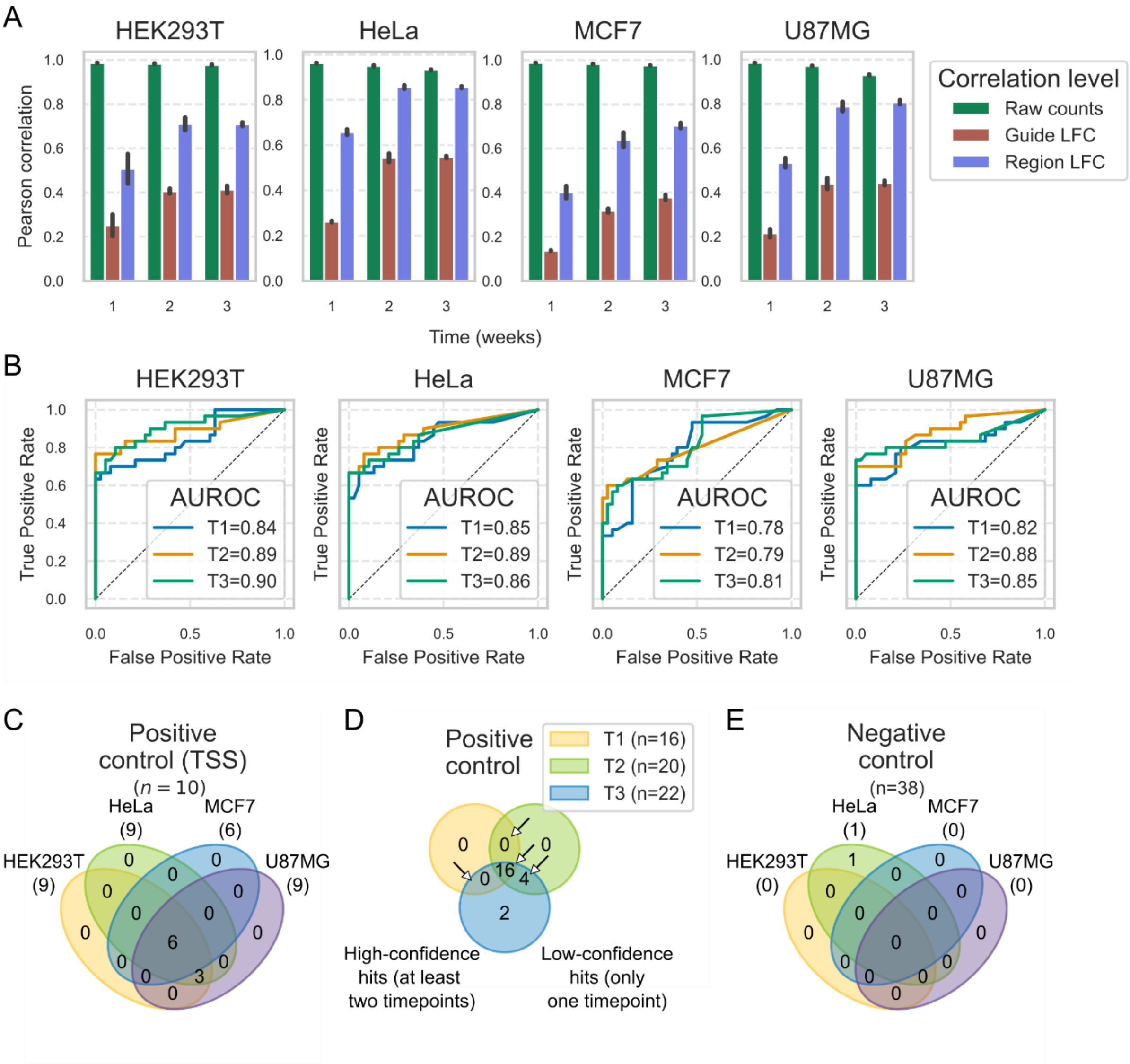


*Figure S2: Results of the paired CRISPR screen. (A) Correlation between CRISPR screen replicates for each cell line. Mean values are plotted with error bars showing standard error. (B) ROC curve for each cell line and timepoint screened, the area under the curve (AUC) is shown in the plot. The dashed line indicates the baseline performance. (C) Number of TSS targeting positive controls identified as hit in the screen. (D) Number of hits in HeLa cells at each timepoint out of the total 30 positive controls. Arrows indicate high confidence hits. (E) Number of negative controls identified as hit in the screen.*

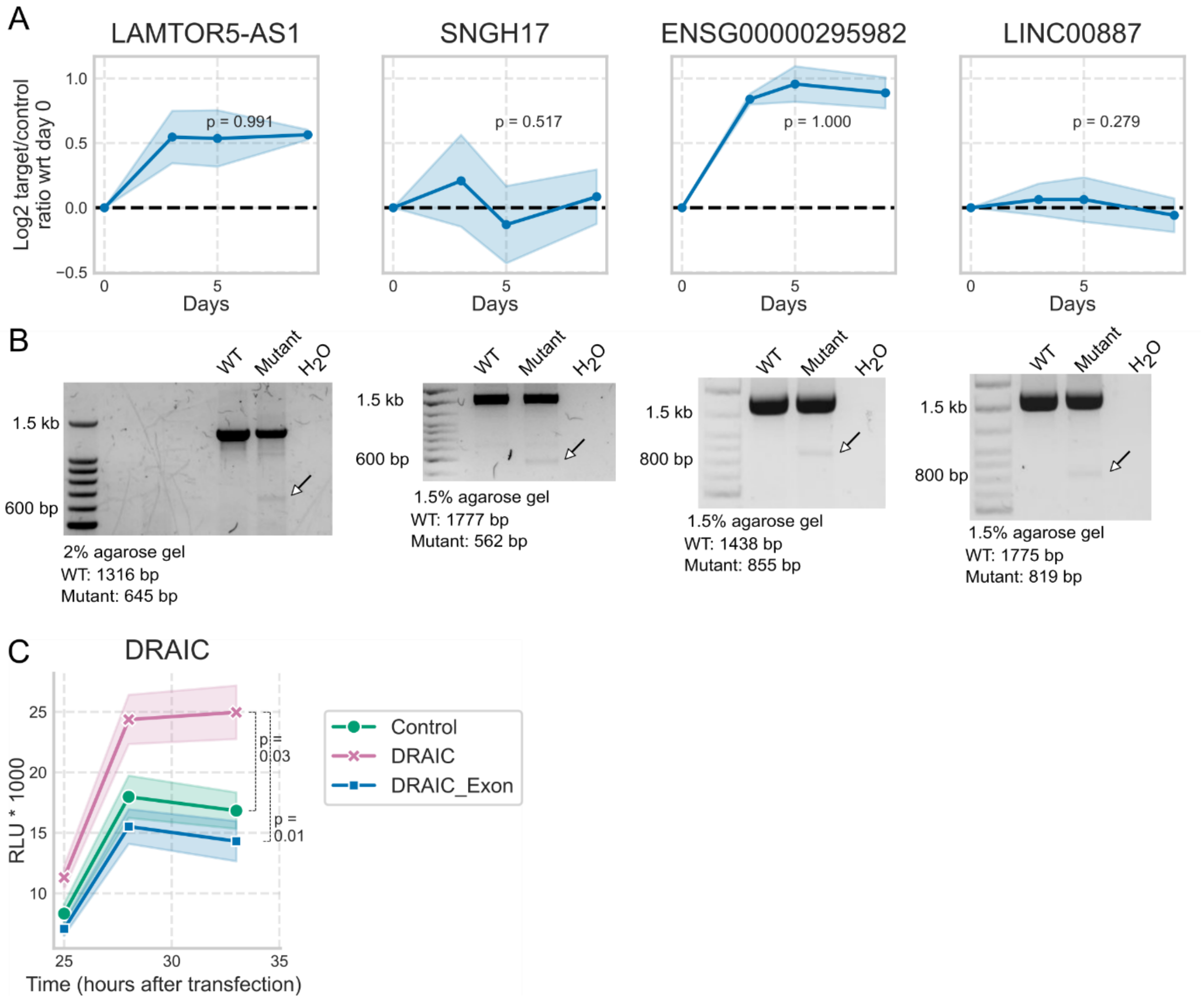


*Figure S3: Validation of the screened exons. (A) Results of fitness competition assay for non-hit exons in the indicated genes. (B) The genomic deletion was confirmed by PCR using primers flanking the deletion region. The arrows indicate the DNA fragment with the expected deletion. (C) Growth curve showing the luminescence values for HeLa cells transfected with different constructs: empty plasmid (control), WT DRAIC or DRAIC Exon 5. Error bars indicate standard error with 3 biological replicates. Statistical significance was calculated using one sided Student's t-test by comparing area under the growth curve.*

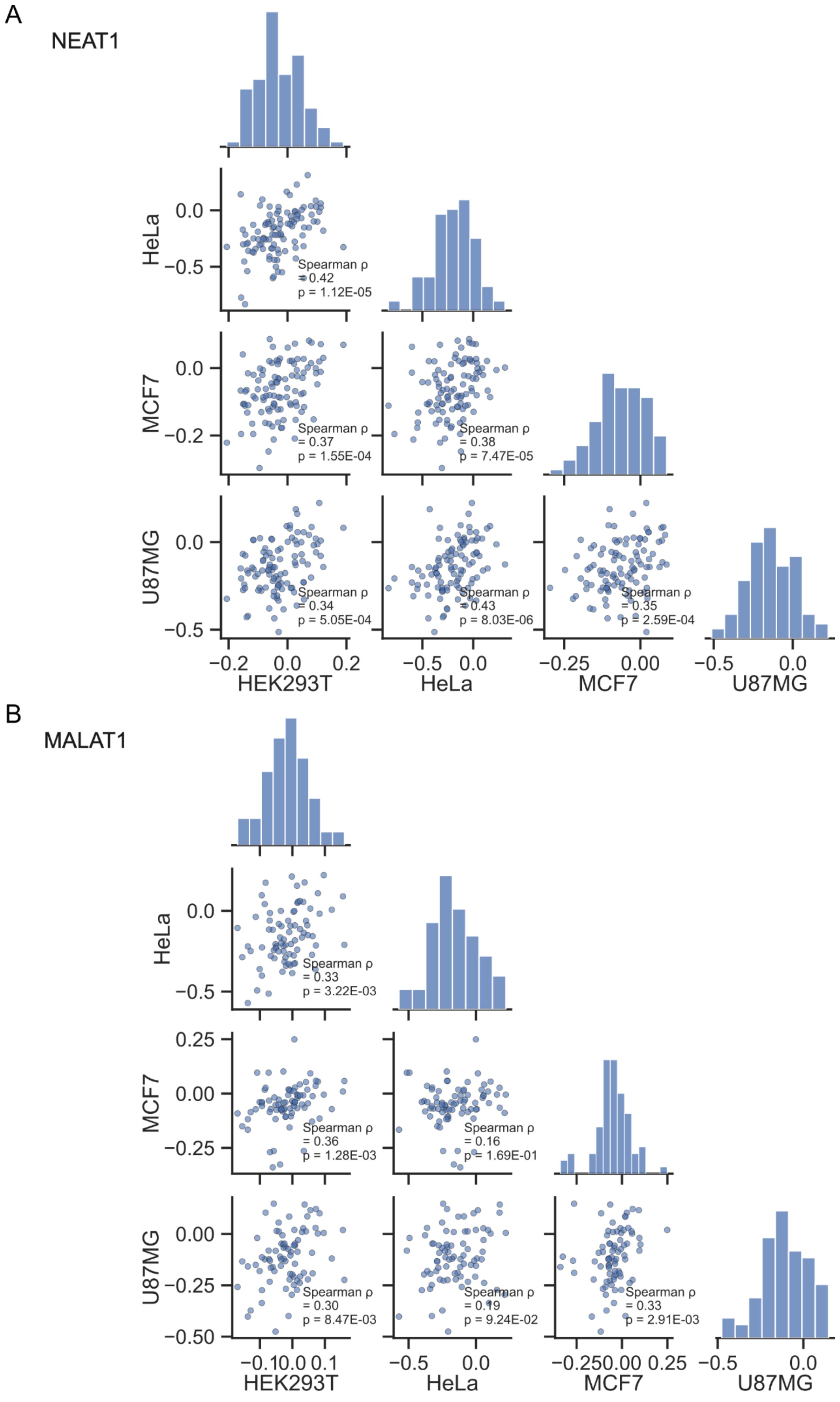
A
NEAT1
HeLa
MCF7
U87MG
HEK293T
0.0
−0.5
−0.2
0.2
−0.25
0.00
Spearman ρ
= 0.42
p = 1.12E-05
Spearman ρ
= 0.37
p = 1.55E-04
Spearman ρ
= 0.38
p = 7.47E-05
Spearman ρ
= 0.34
p = 5.05E-04
Spearman ρ
= 0.43
p = 8.03E-06
Spearman ρ
= 0.35
p = 2.59E-04
B
MALAT1
0.25
−0.25
−0.50
−0.1
0.1
Spearman ρ
= 0.33
p = 3.22E-03
Spearman ρ
= 0.36
p = 1.28E-03
Spearman ρ
= 0.16
p = 1.69E-01
Spearman ρ
= 0.30
p = 8.47E-03
Spearman ρ
= 0.19
p = 9.24E-02
Spearman ρ
= 0.33
p = 2.91E-03

*Figure S4: LFC correlation of all the pgRNAs targeting NEAT1 (A) and MALAT1 (B) for all combinations of cell lines. The Spearman correlation and the corresponding p value is shown inside each scatter plot. The plots on the diagonal represent the distribution of LFC for each cell line.*

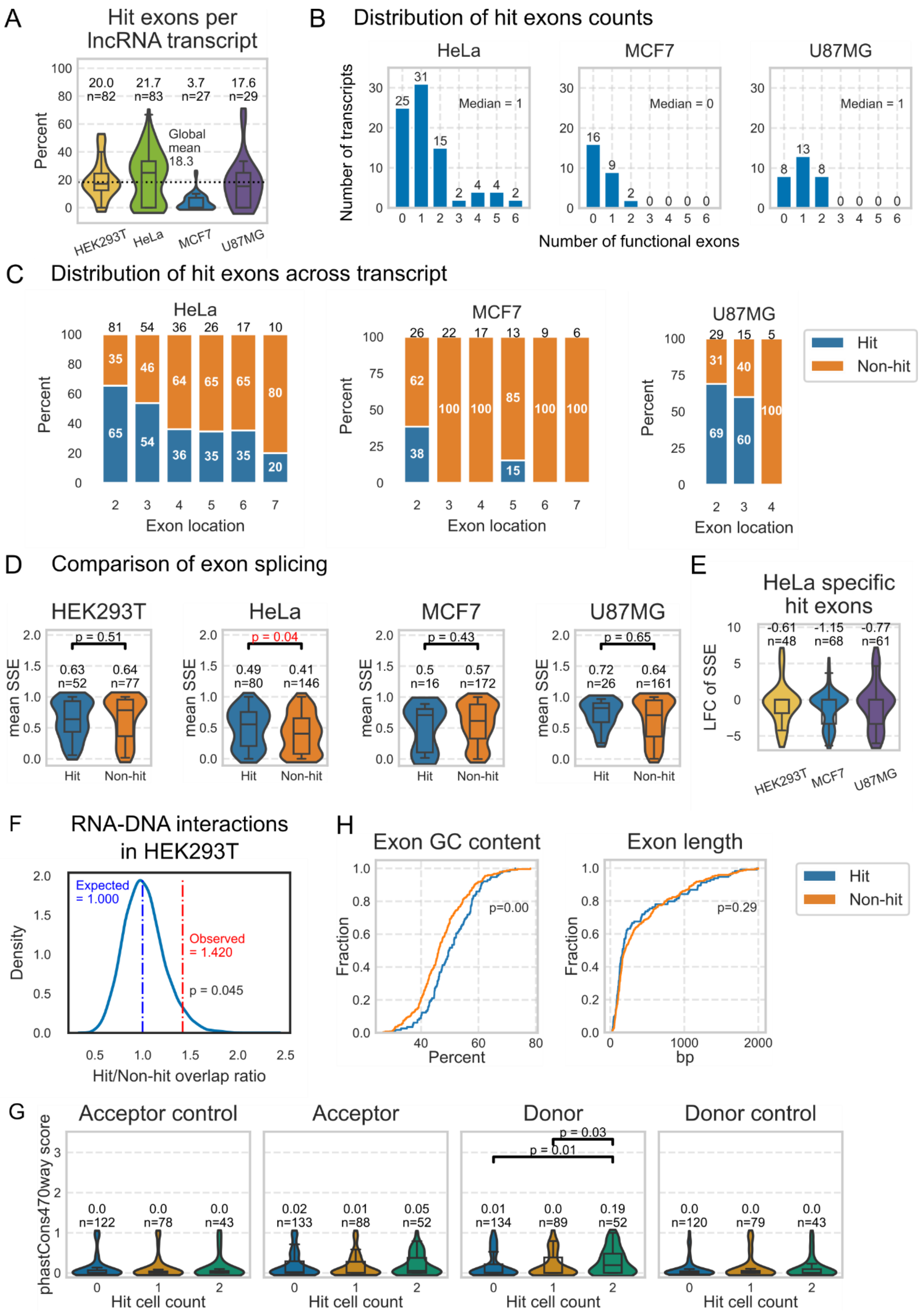


*Figure S5: Properties of lncRNA exons. (A) Distribution of hit exons per transcript in hit lncRNAs, the mean value is shown for each cell line and the number represents the sample size (total number of hit lncRNA transcripts). The global mean across all the cell lines is indicated by the dashed line. (B) Distribution of hit exon counts per transcript in hit lncRNAs. (C) Distribution of hit exon across transcript*

*in hit lncRNAs. The numbers above the bars represent the total number of exons targeted at each location, aggregated from the entire transcript population. (D) Comparison of mean splice-site strength estimate (SSE) of hit and non-hit exons. Statistical significance was calculated using Mann-Whitney U test. (E) Distribution of log2 fold change (LFC) of HeLa specific hit exons' SSE in other cell lines relative to SSE in HeLa. (F) Sampling distribution of ratio of HEK293T hit and non-hit exons overlap with RNA-DNA interaction sites. The expected (median of the distribution) and observed overlap along with p value is mentioned in the plot. (G) PhastCons470way score at the splice sites (labelled as Acceptor and Donor) or matched dinucleotides in the introns (labelled as Acceptor control and Donor control) averaged in a 5bp window surrounding the splice site boundary. Exons are grouped by the number of cell lines in which they are hit: '0' represents non-hits, '1' represents hit only in one cell line, '2' represents hit in at least two cell lines. (H) Cumulative distribution of overlap of exons with GC content and length of hit and non-hit exons aggregated across all cell lines. The p value was calculated using Mann-Whitney U test.*

### Supplementary Files

s1_guides_control.tsv: pgRNAs targeting control regions
s2_guides_lncrna.tsv: pgRNAs targeting lncRNAs regions
s3_mageck_result.tsv: output of mageck analysis
s4_exons_merged.tsv: merged exons
wt1.seq, wt2.seq: Sanger sequencing results of DRAIC transcript before deletion
mutant.seq: Sanger sequencing results of DRAIC transcript after deletion

### Supplementary Table

The IDs of pgRNAs mentioned in these tables correspond to those in the file s1_guides_control.tsv and s2_guides_lncrna.tsv

Supplementary Table 1

| Description | Sequence |
|---|---|
| Fig 1C (gRNA sequence targeting MALAT1 enhancer) | gRNA1: GTTGGTCAAGTAAAGACACG<br>gRNA2: AGTTCTGCCTCAGCTCAGGA |
| Fig 1C (gRNA sequence targeting GFP) | Addgene, #78535<br>gRNA1: GAGCTGGACGGCGACGTAAA<br>gRNA2: CAGAACACCCCCATCGGCGA |
| Fig 1C (qPCR primers) | GAPDH_F: GCACCGTCAAGGCTGAGAAC<br>GAPDH_R: TGGTGAAGACGCCAGTGGA<br>MALAT1_F: GCTGGGGAATCCACAGAGAC<br>MALAT1_R: CATCTCAGCCCTTGTTATCCTG |
| Fig 1C (PCR primers for gel electrophoresis of amplicons) | MALAT1_F: CCTGCTATGAACTGACCCATG<br>MALAT1_R: CCTGAACAGTCAGTCCATGCT |

Supplementary Table 2

| Description | Sequence |
|---|---|
| Fig 3D (DNA primers) | DRAIC_F: GACATTGTGCTGGGGAAGGA |

| | |
|---|---|
| | DRAIC_R: TGCAGTGACCCCATTAGCAG |
| Fig 3D (RNA primers, 1st RT-PCR followed by a nested PCR) | DRAIC_F1: CATTCTCCTGCCTCAGCCTC<br>DRAIC_R1: TAGGTGCGTTGGCTATGTCC<br>DRAIC_F2: ACAATTTCACTGAAGTATTGCTTGT<br>DRAIC_R2 TGCTTTTCGTTGTTGAGCGT: |

Supplementary Table 3

| Description | Sequence / pgRNA ID |
|---|---|
| Fig 4A (AAVS1 GFP) | Addgene, #141211<br>gRNA1: GACCTGCTCACAGGCGAGGTA<br>gRNA2: ACCTGGTTCACACGGCGCAG |
| Fig 4B (RPS5 mCherry) | gRNA1: GACCTGCTCACAGGCGAGGTA<br>gRNA2: ACCTGGTTCACACGGCGCAG |
| Fig 4B (AAVS1 mCherry) | gRNA1: GAGTGCCCTTGCTGTGCCGC<br>gRNA2: CCTCTGGGGGATGCAGGGGA |
| Fig 4C (DRAIC) | Exon 5: pgRNA6039<br>Exon 6: pgRNA6052_1 and pgRNA6053_2 |
| Fig 4D (LINC01806) | Exon 2: pgRNA911<br>Exon 3: pgRNA7965 |
| Fig S3A (LAMTOR5-AS1) | pgRNA13 |
| Fig S3A (SNHG17) | pgRNA3752 |
| Fig S3A (ENSG00000295982) | pgRNA93 |
| Fig S3A (LINC00887) | pgRNA3822 |

Supplementary Table 4

| Gene | Transcript | Exon |
|---|---|---|
| DRAIC | ENST00000558633.6 | 4 and 5 |
| EWSAT1 | ENST00000558369.5 | 3 |
| LINC01270 | ENST00000653970.1 | 2 |
| PVT1 | ENST00000657449.1 | 11 |

Supplementary Table 5

| Gene | Element (hg38 coordinates) | Scramble |
|---|---|---|
| DRAIC (ENST00000558633.6) | SINE repeat (chr15:69468334-69468393) | GTCGAGCTAGTAATCCTATTTTTTT<br>TGCGACTCGCAGCTCGTAGTTTG<br>C<br>TTCGCCCTC |
| LINC01270 (ENST00000653970.1) | ILF3 motif (chr20:50293526 - 50293533) | TAGCGTC |
| LAMTOR5-AS1 (ENST00000608486.5) | U2AF2 CLIP peak (chr1: 110412172-110412208) | GCAAATTCCTCTGCGGTCACTCG<br>TCACGTATAAGA |

| | | |
|---|---|---|
| LAMTOR5-AS1 (ENST00000608486.5) | PhastCons (chr1: 110412246-110412253) | TATCGTG |
